# Multivalent peptide ligands to probe the chromocenter microenvironment in living cells

**DOI:** 10.1101/2021.11.15.468638

**Authors:** Nora Guidotti, Ádám Eördögh, Maxime Mivelaz, Pablo Rivera-Fuentes, Beat Fierz

**Author notes:** These authors contributed equally (Nora Guidotti, Ádám Eördögh).

## Abstract

Chromatin is spatially organized into functional states that are defined by both the presence of specific histone post-translational modifications (PTMs) and a defined set of chromatin-associated ‘reader’ proteins. Different models for the underlying mechanism of such compartmentalization have been proposed, including liquid-liquid phase separation (LLPS) of chromatin-associated proteins to drive spatial organization. Heterochromatin, characterized by lysine 9 methylation on histone H3 (H3K9me3) and the presence of heterochromatin protein 1 (HP1) as a multivalent reader, represents a prime example of a spatially defined chromatin state. Heterochromatin foci exhibit features of protein condensates driven by LLPS; however, the exact nature of the physicochemical environment within heterochromatin in different cell types is not completely understood. Here, we present tools to interrogate the environment of chromatin sub-compartments in the form of modular, cell-permeable, multivalent and fluorescent peptide probes. These probes can be tuned to target specific chromatin states by providing binding sites to reader proteins and can thereby integrate into the PTM-reader interaction network. As a target, here we generate probes specific to HP1, directing them to heterochromatin at chromocenters in mouse fibroblasts. Moreover, we use a polarity-sensing photoactivatable probe that photoconverts to a fluorescent state in phase-separated protein droplets and thereby reports on the local microenvironment. Equipped with this dye, our probes indeed turn fluorescent in murine chromocenters. However, image analysis and single-molecule tracking experiments reveal that the compartments are less dense and more dynamic than HP1 condensates obtained *in vitro*. Our results thus demonstrate that the local organization of heterochromatin in chromocenters is internally more complex than an HP1 condensate.

## Introduction

The eukaryotic genome is packed into chromatin, which adopts a folded structure and governs DNA accessibility^1^. Nucleosomes form the fundamental chromatin building blocks and contain around 147 bp of DNA wrapped around a protein core composed of histones H2A, H2B, H3, and H4^2,3^. Histones are heavily modified by combinations of post-translational modifications (PTMs) that epigenetically control essential regulatory processes, e. g. gene expression and repression. PTMs recruit specific effector proteins^4^ that, in turn, organize the chromatin fiber into three-dimensional sub-nuclear compartments and define complex chromatin states, including relaxed and active euchromatin or compacted and transcriptionally silenced heterochromatin. In particular in murine cells, extended regions of heterochromatin at pericentric satellite repeats form highly visible compact compartments cells called chromocenters^5^.

Heterochromatin silencing relies on the recruitment of HP1 proteins, which in mammalian cells include paralogs HP1α, HP1β, or HP1γ^6^. HP1 proteins specifically bind to histone H3 di- and tri-methylated at lysine 9 (H3K9me2 and me3) via their chromodomain (CD)^7,8^. A flexible and unstructured hinge region connects the HP1 CD to a dimerization motif, the chromoshadow domain (CSD). Additionally, the CSD acts as a platform for recruiting other repressive effectors, thus contributing to heterochromatin spreading along the genome^9,10^. Structurally, HP1 dimerization and oligomerization^11,12^ enables multivalent chromatin recognition^13^. Association of HP1 dimers and oligomers to H3K9me3 on different nucleosomes results in cross-bridging of chromatin strands^11,14-17^. Such chromatin bridging can result in chromatin compaction and the formation of compact chromatin ‘globules’^18^.

Recent experiments further identified liquid-liquid phase separation (LLPS) of HP1 proteins as a major contributor to heterochromatin organization^12,19^. Indeed, LLPS has been proposed to represent a general mechanism for genome regulation^20^. LLPS is driven by multivalent weak interactions between macromolecules and often involves intrinsically disordered protein regions^21^. Phase-separated condensates selectively accumulate specific regulatory factors in a highly dynamic way while non-interacting macromolecules are excluded^22^. LLPS thus represents an attractive model for nuclear compartmentalization and spatiotemporal control of biochemical reactions in cell nuclei^20^. In the process of heterochromatin establishment, phase-separated HP1 droplets are thought to envelop heterochromatin regions establishing a sub-compartment separated from the nucleoplasm by a boundary that restricts biochemical access^12,19,23^.

The globule and the LLPS models for heterochromatin formation are not mutually exclusive and may depend on cell type and developmental stage^19,23^, but a general model of heterochromatin compaction remains elusive. Therefore, the investigation of the biophysical, biochemical, and structural principles underlying heterochromatin organization, e. g. via biomolecular condensates or through the establishment of a cross-bridged globular state is key to understanding cell function.

We recently developed genetically-encoded multivalent reader proteins coupled to fluorescent proteins to image chromatin states in living cells^24^. These probes are characterized by defined combinations of histone PTMs. Similar multivalent readers have been used to characterize the chromatin PTM-dependent interactome at given chromatin states^25^. To image distinct yet diverse chromatin states in cells, larger modularity and decreased complexity of the probes would be advantageous. In contrast to proteins, peptides are smaller and can be delivered efficiently to cells using appropriate cell-penetrating sequences^26^. Here, we report a modular strategy to synthesize cell-permeable, multivalent, peptide-based probes that selectively accumulate in the nucleus of live cells at specific chromatin compartments. We targeted heterochromatin compartments by exploiting binding interactions to HP1 proteins. Using these novel reagents, coupled to fluorescent dyes and environmentally sensitive probes, we were able to visualize the organization of constitutive heterochromatin in live cells and probe their environment. Our results reveal that chromocenters in mouse fibroblast cells are of lower density and higher polarity than nucleoli and phase-separated HP1α droplets prepared *in vitro*. Thus, in this cell type, the internal environment of chromocenters is more complex than expected if the compartment were purely defined by HP1 phase separation.

## Results

### Designing multivalent peptide ligands for chromatin readers

We set out to develop modular peptide reagents to target, visualize and probe a defined chromatin state characterized by the presence of distinct combinations of chromatin readers and histone PTMs in living cells. We reasoned that multivalent peptide ligands could serve as a starting point for successful designs. We further decided to use histone-derived peptides carrying defined PTMs for targeting, as they interact with high specificity with accumulated reader proteins present at a given chromatin state. As an initial target, we selected chromocenters, foci of dense HP1-based heterochromatin at silenced pericentric satellite repeats^5^, allowing us to probe their microenvironment (**Figure 1A**).

**Figure 1.**
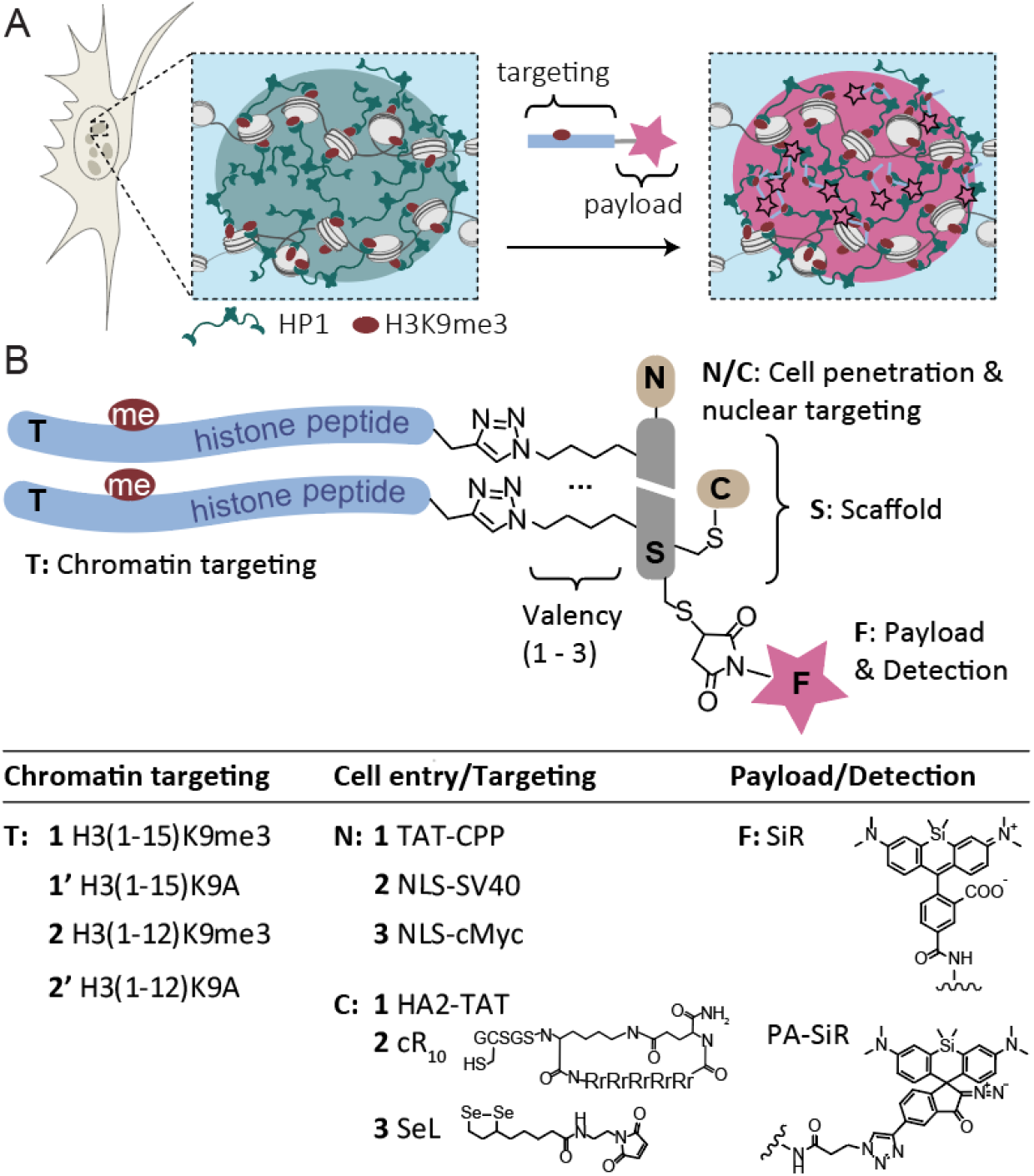
Peptide reagents to probe chromocenters. **A)** Scheme of a chromocenter containing heterochromatin established via nucleosome cross-bridging by HP1 proteins that bind H3K9me3. Moreover, HP1 proteins may phase-separate to form HP1 droplets surrounding the heterochromatin focus. Peptides containing an HP1 binding site can invade the sub-compartment and deliver a payload (i. e. a dye) for imaging of the chromatin state. **B)** General design of the peptide reagents containing a scaffold (**S**) to which chromatin targeting peptides (**T**), cell-penetrating- or nucleus-targeting moieties (**N/C**), and a fluorescent payload (**F**) can be attached. A table of the explored combinations is provided below. For all synthesized peptides and probes, see **Tables S1, S2**.

We identified a number of challenges that had to be overcome: the reagent needed to (i) penetrate the cell membrane; (ii) enter and stay in the nucleus; and (iii) specifically bind to the reader protein. To address these challenges, we settled on a modular design principle. A central peptide scaffold (**S**) was equipped with two or three attachment points for targeting peptides (**T**), introduced via copper-catalyzed azide-alkyne Huisgen cycloaddition (CuAAC)^27^ (**Figure 1B**). As for targeting peptides (**T1, T2**), we chose sequences derived from the H3 tail containing a trimethylated lysine (equivalent to K9 in H3) surrounded by its native sequence, resulting in an HP1 binding site^7,8,28^. We further included H3 peptides where the critical lysine was mutated to alanine (K9A) as non-binding controls to test target specificity (**T1’, T2’**).

The scaffold peptide was designed to contain further points for derivatization: To optimize cell and nuclear entry, we explored the addition of a range of cell-entry and nuclear targeting moieties. At the scaffold N-terminus, additions included a cell-penetrating peptide (CPP) derived from the HIV TAT protein^29,30^ (TAT-CPP, **N1**), a nuclear localization sequence (NLS) derived from the SV40 T antigen^31^ (NLS-SV40, **N2**) or an NLS derived from the human c-Myc protein^32^ (NLS-cMyc, **N3**). To improve cell-entry further, we also explored attaching a pH-sensitive endosomolytic peptide derived from the HA2 subunit of the influenza viral hemagglutinin protein^33^ (HA2-TAT, **C1**), an optimized CPP composed of a cyclic polyarginine sequence^34^ (cR10, **C2**), and a diselenolane compound to improve cellular uptake^35^ (SeL, **C3**). Finally, a ‘payload’ was attached in the form of a far-red dye (silicon rhodamine, **SiR**^36^) to image heterochromatin loci or a polarity-sensing photoactivatable dye^37^ (**PA-SiR**) to probe the heterochromatin environment.

For evaluation and optimization of the different designs, all peptides (scaffolds, targeting moieties, CPPs, NLS carrying peptides) were individually synthesized by solid-phase peptide synthesis (Fmoc-SPPS) and assembled in a modular fashion (see **Supplementary Figures S1 – S3** for peptide synthesis). The key reaction in probe assembly, which remained constant throughout all designs, was the attachment of the H3-derived targeting peptides to a dye-derivatized scaffold (**Figure 2A**). Using probe **(T2)**_**3**_**-SN3-SiR** as an example, we synthesized and purified a scaffold peptide carrying the cMyc NLS (**N3**), three azido-lysines flanked by flexible linker residues (glycine-serine), an S-acetamidomethyl (ACM)-protected cysteine as well as a free cysteine residue (**SN3, Figure 1A, Figure S1**, and **Table S1**).

**Figure 2:**
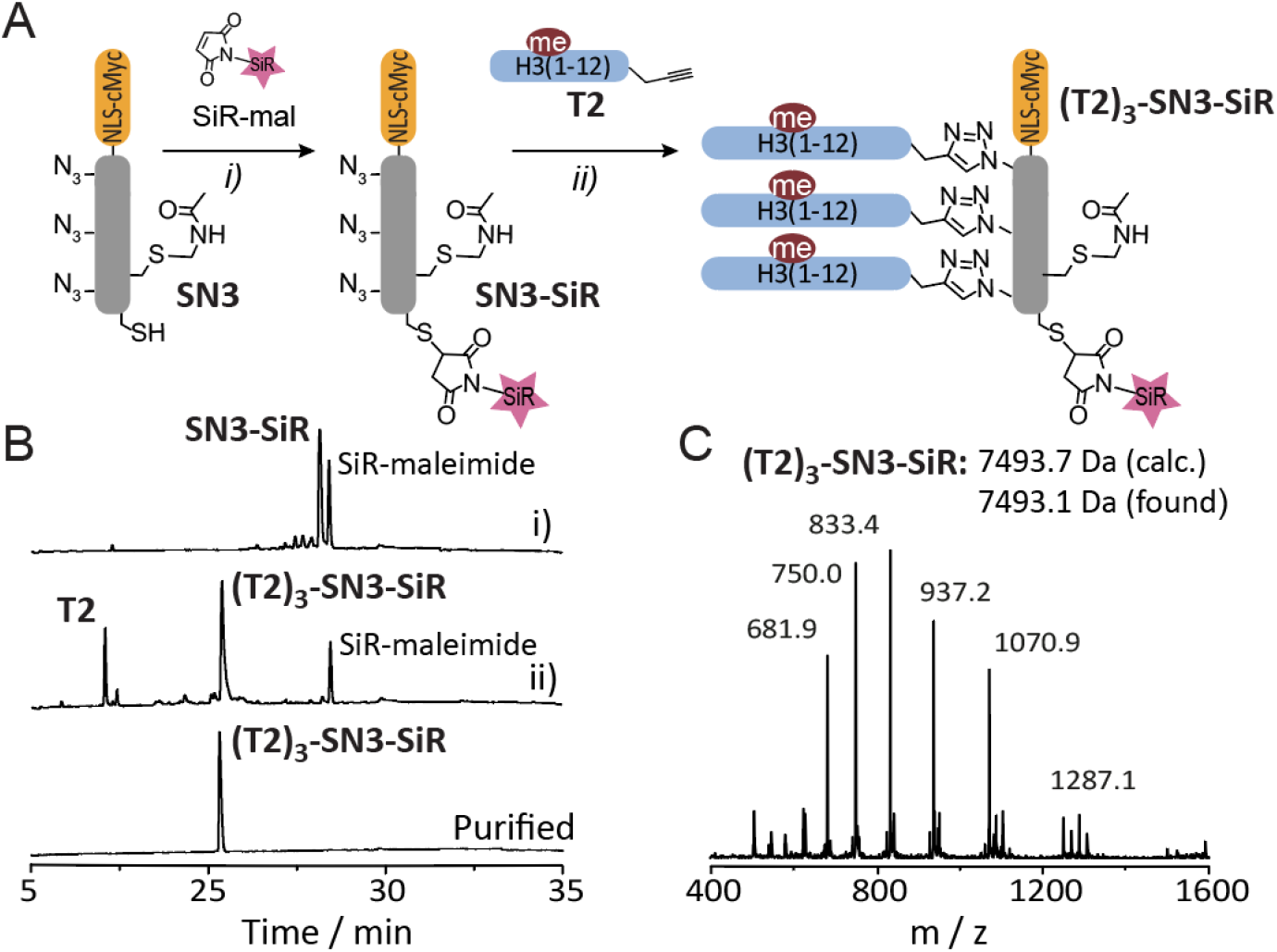
Probe synthesis. **A**) Scheme of synthesis of probe **(T2)**_**3**_**-SN3-SiR** in a two-step, one-pot procedure: i) coupling of SiR-maleimide followed by attachment of the targeting peptides **T2** via CuAAC. **B**) HPLC analysis of reaction progress: i) SiR-mal coupling (t = 15 min), ii) CuAAC of **T2** (t = 5 min), and final purified product. **C**) ESI-MS analysis of **(T2)**_**3**_**-SN3-SiR** showing the expected molecular mass (7439.7 Da, calculated, 7439.1 Da measured). Further analysis of targeting and cell-penetrating peptides can be found in **Figures S2, S3**.

Treatment of the scaffold peptide **SN3** with SiR-maleimide generated the SiR-modified scaffold peptide (**SN3-SiR**) as the sole product (**Figure 2B**). This peptide was further functionalized with histone tail peptide **T2** under standard CuAAC conditions to give probe peptide (**T2)**_**3**_**-SN3-SiR** in good overall yields (see supporting information for experimental details). In the case where further functionalization of the probe was needed (i.e., coupling of a cell-delivery agent), the acetamidomethyl (Acm) group could be chemoselectively removed, and the thiol reacted with the desired moiety. Exploiting the highly modular nature of our strategy, we were able to couple different scaffold peptides to various histone fragments in a combinatorial way (see **Tables S1, S2**).

### Optimization of chromocenter-targeting peptide probes

Having established a general probe design and synthesis procedures, we started to test different designs by evaluating their ability to enter cells and co-localize with chromocenters. As chromocenters are especially prominent in mouse cells ^38^, we transfected NIH-3T3 murine embryonic fibroblasts with HP1α fused to the fluorescent protein mEOs3.2^13^ for direct quantification of targeting. We then treated the cells with peptide probes (using concentrations between 1 and 10 μM) for 1 h, followed by medium exchange and imaging via confocal microscopy. We employed either the HP1α-mEOs3.2 signal or a DNA stain (Nuclear-green) to mark chromocenters and the far-red channel to image the peptide probes.

We assessed the ability of the probes to enter cells and target chromocenters, based on various properties, including the nature of cell-penetrating and nuclear targeting moieties, the valency of the scaffold, and the chromatin targeting peptides. Initially, we evaluated a bivalent peptide design, **(T1)**_**2**_**-SN1-SiR**, carrying the TAT-CPP for cell entry and H3(1-15)K9me3 as targeting peptide (see **Figure S4** for synthesis of probes). Whereas the peptide efficiently entered the cell, it did not reach the nucleus but accumulated in cytoplasmic vesicles (**Figure 3A**). We thus combined the probe **(T1)**_**2**_**-SN1-SiR** with the endosomolytic peptide HA2-TAT (**C1**), which has been used to deliver cargo to nuclei^39^. Co-incubation of **(T1)**_**2**_**-SN1-SiR** with HA2-TAT at an equimolar concentration significantly improved cellular uptake and nuclear localization of the probe (**Figure 3B**). However, whereas TAT-CPP aided cell entry and served as a nuclear import tag, the probe accumulated in nucleoli, potentially due to the positively charged nature of the TAT-CPP sequence (**Table S3**). Conversely, probe **(T1)**_**2**_**-S-SiR**, lacking this CPP, entered cells in the presence of HA2-TAT but was again excluded from the nucleus (**Figure 3C**), demonstrating the need for a nuclear import signal.

**Figure 3:**
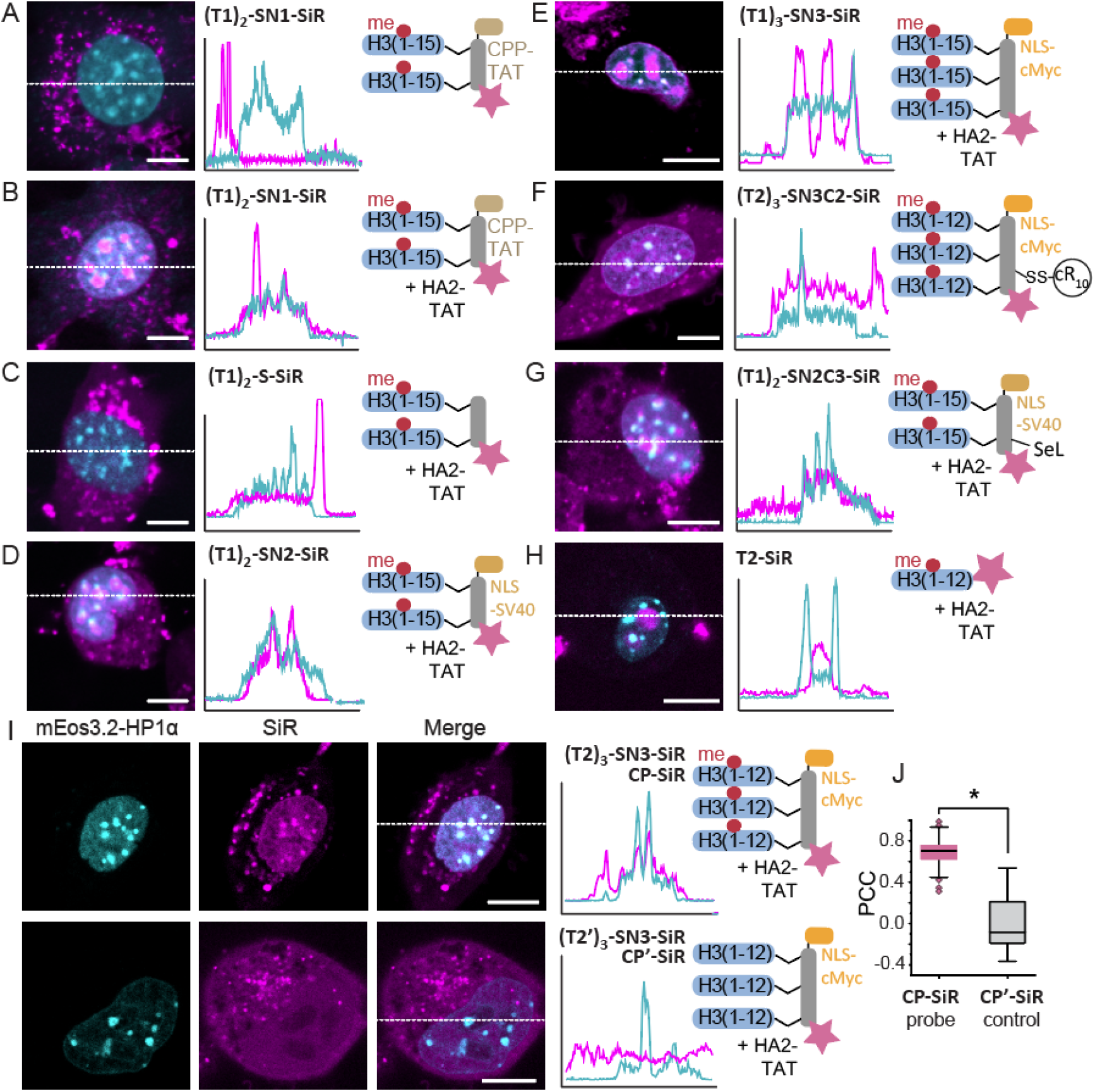
Development of a heterochromatin targeting peptide probe. **A-H**) Confocal microscopy images of 3T3 fibroblasts treated with the indicated peptides. SiR fluorescence emission of the indicated peptide probes (magenta) is overlaid with HP1α-mEOs3.2 fluorescence (cyan). 2D intensity profiles corresponding to the dotted lines displayed on the images (line scans) are depicted to the right of the micrographs. **I**) Confocal microscopy images and line scans of 3T3 fibroblasts treated with **CP-SiR** probe (i.e. **(T2)**_**3**_**-SN3-SiR**, top) or the K9A variant **CP’-SiR. J**) Correlation between HP1α-mEOs3.2 and **CP-SiR** or **CP’-SiR** staining, demonstrating good colocalization for **CP-SiR**, and the dependence on the presence of the H3K9me3 modification in **CP-SiR** for heterochromatin staining (see Supplementary Materials for details on the calculation). N = 21 cells (CP-SiR), N = 10 cells (CP’-SiR). PCC: Pearson correlation coefficient. Two-sample t-test *p<4·10^−7^, Scale bars: 10 μm.

Consequently, we replaced TAT with the NLS-SV40 sequence^31^ (**N2**), which enabled efficient cell penetration and nuclear import in the presence of HA2-TAT (**Figure 3D**), but the **(T1)**_**2**_**-SN2-SiR** probe still accumulated preferentially in nucleoli. We thus exchanged NLS-SV40 for NLS-cMyc, which is substantially less polar^32^. Moreover, we increased multivalency to n = 3 to further direct the probes to chromocenters. The resulting probe **(T1)**_**3**_**-SN3-SiR** again entered cells and nuclei when co-administered with HA2-TAT but showed intense nucleolar staining (**Figure 3E**). In parallel, we also tested cell delivery via alternatives to HA2-TAT, in particular using the cyclic arginine peptide cR10 (**Figure S5**) or diselenolane reagents (**Figure S6**). In both cases, cell entry was very efficient (**Figure 3F, G**), but chromocenter targeting was not yet ideal. As a control, we also tested the monovalent peptide, comprised solely of H3(1-12)K9me3-SiR (**T3-SiR**), which did not show any chromocenters localization, demonstrating that multivalency is critical to reaching heterochromatin loci (**Figure 3H**).

Finally, the best results were obtained with the trivalent peptide **(T2)**_**3**_**-SN3-SiR** using a shortened H3 sequence omitting K14, which exhibited high cell permeability in the presence of HA2-TAT, low nucleolar staining, and selectively accumulated in chromocenters (**Figure 3I, J**). Importantly, chromocenters accumulation was directly dependent on the presence of K9me3, as the control probe **(T2’)**_**3**_**-SN3-SiR**, carrying an alanine instead of lysine 9, did not show any colocalization (**Figure 3I, J**). In conclusion, by optimizing our modular system, we engineered the peptide reagent **(T2)**_**3**_**-SN3-SiR** that efficiently targets HP1 foci in living cells. From here on, we refer to this construct as **CP-SiR** (for chromocenter probe labeled with SiR).

### A polarity sensor for HP1α condensates

The chromocenter-directed peptide probe **CP-SiR** is useful to image subnuclear chromatin states. In addition, however, this probe can also be equipped with a sensor moiety to probe the physicochemical environment in a given nuclear compartment. The local structure and properties of chromocenters in specific cells, in particular regarding HP1 phase separation, is a subject of active research^12,23^. We posited that **CP-SiR** and derivatives could be used to gain insight into the environment within chromocenters. The photoactivatable dye PA-SiR (**Figure 1B**) contains a diazoindanone moiety, spirofused to a silicon-containing rhodamine dye, and is non-fluorescent^37,40^. Upon irradiation, it undergoes a photoinduced Wolff rearrangement, which follows divergent pathways dictated by the polarity of the environment^37^. In an apolar environment, PA-SiR photoconverts into a strongly fluorescent molecule with spectral characteristics that are nearly identical to those of SiR. In contrast, in a highly polar environment, i.e., in an aqueous solution, the major rearrangement product is a ring-expanded, non-fluorescent product (**Figure S7**). Notably, the fraction of fluorescent photoproduct directly reflects the overall polarity of the medium, as revealed by systematic experiments under different solvent conditions^37^. We thus decided to use this property of PA-SiR to investigate the polarity of the environment within chromocenters. Moreover, the photochemistry of PA-SiR allows for sparse activation of molecules suitable for single-molecule tracking (SMT). We envisioned that these tracking experiments could be used to estimate the diffusion coefficients of molecules within chromocenters.

To validate these ideas, we first investigated the ability of **CP-SiR** to partition into HP1α condensates *in vitro*. We expressed HP1α in the presence of casein kinase 2 (CK2), resulting in the phosphorylated form of HP1α (pHP1α)^41^, which has a strong propensity to phase-separate at micromolar concentrations^12^. The addition of crowding agents, such as polyethylene glycol (PEG), lowers the concentration threshold for such phase transition^12^. Indeed, the formation of pHP1α droplets was induced from a solution of 40 μM pHP1α in the presence of 7.5% PEG (**Figure 4A**). Added **CP-SiR** partitioned into pHP1α phase, as revealed by an increase in fluorescence emission in the droplets and a reduction in background fluorescence from the surrounding medium (**Figure 4B**, final concentrations: 30 μM pHP1α, 5% PEG, 100 nM **CP-SiR**). In contrast, the control peptide carrying a K9A substitution (**CP’-SiR**) did not strongly partition into the pHP1α phase, demonstrating specific targeting.

**Figure 4:**
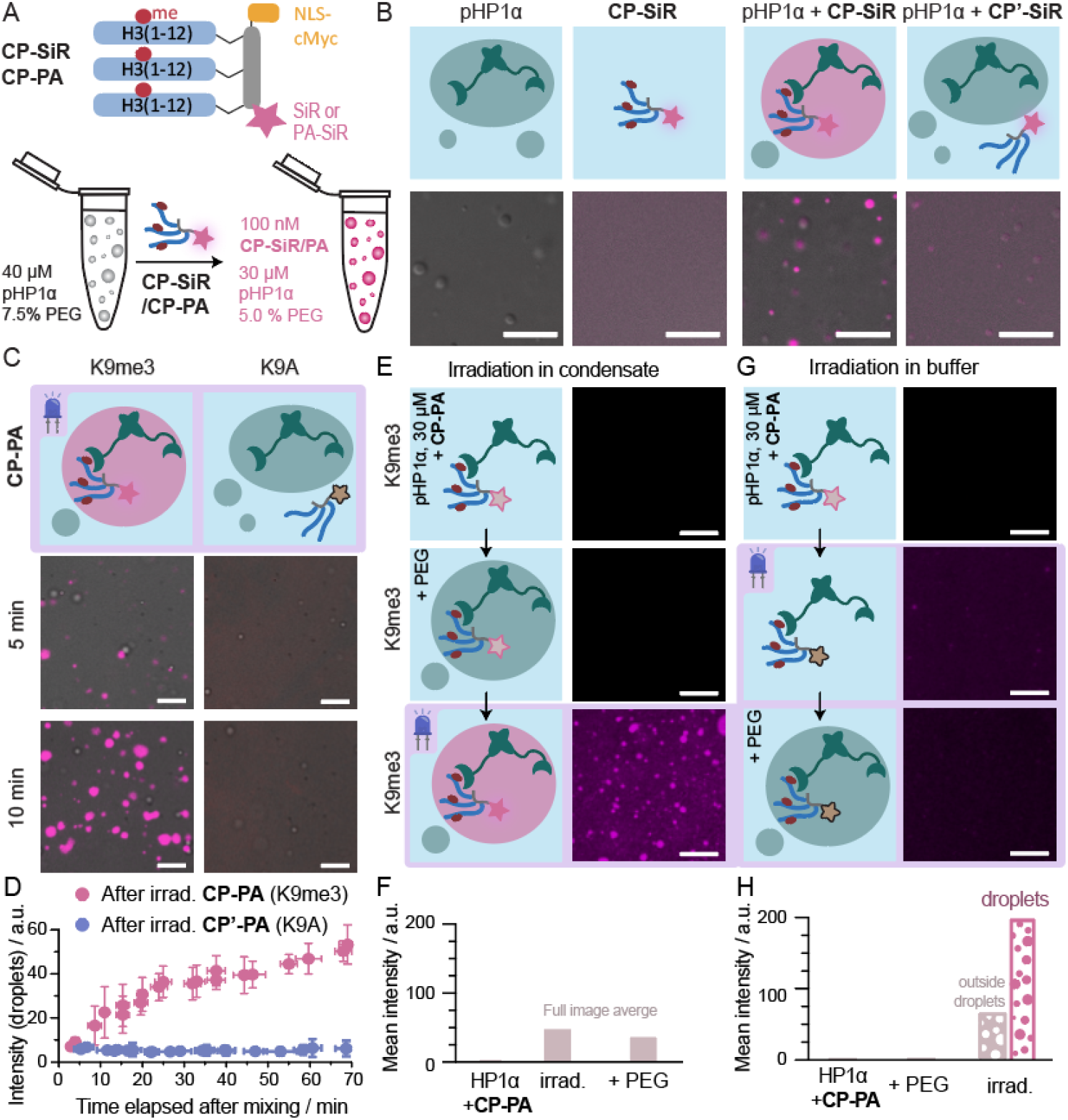
CP-PA is specifically photoactivated in pHP1α droplets. **A)** Scheme of the partitioning experiments (for details, see text). **B**) Only **CP-SiR**, but not **CP’-SiR**, efficiently partitions into pHP1α droplets. **C**) Phase-separated droplets were generated by mixing pHP1α and PEG at indicated concentrations, then **CP-PA** or control **CP’-PA** was added to the phase-separated sample (t = 0 min) to reach final concentrations pHP1α (30 μM), PEG (5 %_w_), and **CP-PA** (100 nM). The sample was imaged by a confocal microscope by first illuminating the sample with activating laser (405 nm, 200 ms, 65 mW) then reading out the activated signal with (638 nm, 800 ms, 100 mW). For a later timepoint and further characterization of the droplets, see **Figure S8. D**) Quantification of signal intensity from panel **C**. Images were segmented into droplets and background. The intensity measured in droplets is shown, from independent experiments (N=3), error bars represent standard deviation. **E**) **CP-PA**, pHP1α and PEG were mixed and exhaustive irradiation was performed (LED, 405 nm, 2 mW cm^-2^, 150 s) after 10 min. **G**) Quantification of photoactivation of **CP-PA** in panel **E. G**) **CP-PA** was mixed with pHP1α, and the sample was irradiated (LED, 405 nm, 2 mW cm^-2^, 150 s) to convert all photoactivatable **CP-PA** precursor. PEG was added to the system after irradiation. **H**) Quantification of photoactivation of **CP-PA** in panel **G**. Fluorescence turn-on is only observed if irradiation happens in the phase-separated condensate. All scale bars: 10 μm.

We then synthesized another probe containing the environment sensitive PA-SiR dye and bearing the K9me3 modification (**CP-PA**), as well as a K9A containing control probe (**CP’-PA, Figure S7**). When added to pre-made pHP1α droplets, **CP-PA** partitioned into the pHP1α phase and turned fluorescent upon irradiation (**Figure 4C**). Longer incubation times prior to irradiation led to stronger fluorescence signal due to increased partitioning of **CP-PA** into phase-separated droplets (**Figure 4D, Figure S8A**,**B**). In contrast, **CP’-PA** did not partition into droplets and only produced a low amount of fluorescencent product upon irradiation, because most of the precursor was converted to a non-fluorescent isomer in the aqueous phase (**Figure 4C-D, Figure S8C**). During the experiment, pHP1α condensate size increased with sedimentation and coalescence, but the size and shape of the droplets was not significantly influenced by either **CP-PA** or **CP’-PA** probes (**Figure S8E-G**).

The previous experiments provide evidence that, when photoactivatable probes are irradiated in the condensed phase, the bright photoproduct is formed preferentially. However, it was unclear whether the change in photochemical selectivity was caused by the environment in the condensate, or by probe binding to pHP1α proteins. We therefore performed photoactivation of the probe in presence of pHP1α protein in two separate experiments, with and without phase separation. First, we mixed 100 nM **CP-PA** with 30 μM of pHP1α, then added PEG to initiate condensate formation and finally irradiated the sample to photoconvert **CP-PA**. This sequence of events resulted in the generation of fluorescent probe in droplets (**Figure 4E, G**), similar to previous experiments. Second, we incubated **CP-PA** and pHP1α in the absence of PEG, conditions that do not promote the formation of pHP1α condensates. Under these conditions, the peptide was fully bound to the protein^13,41^. However, irradiation with blue light did not result in photoconversion to a fluorescent state, and pHP1α droplets formed after PEG addition did not show significant fluorescence (**Figure 4E, G**). This result demonstrates that pHP1α binding is not sufficient for **CP-PA** photoconversion to a fluorescent state, but the dense and low-polarity environment within pHP1α droplets is necessary.

Having the possibility of selectively turning on **CP-PA** in pHP1α condensates finally allowed us to use SMT to investigate probe dynamics within this phase. Using a low power setting for the activation laser, we activated a small subset of probes and tracked the molecules over 9 min at an 18 Hz framerate. Surprisingly, the overwhelming majority of tracked molecules in pHP1α condensates were essentially immobile (average diffusion coefficient D_average_ = (8.9 ± 11) · 10^−4^ μm^2^ s^-1^ (mean ± CI_95%_, N=4), **Figure S8J**). Individual H3K9me3-carrying peptides interact with pHP1α with a low micromolar *K*_D_^13,41^, which should allow for rapid dissociation-association reactions. However, multivalent interaction could strongly decrease probe mobility. Moreover, upon photoactivation, **CP-PA** reacts with a nucleophile to transition to a fluorescent state. In condensates, the high pHP1α density increases the probability of the probe crosslinking to nearby proteins, which contributes to the observed low diffusion coefficient.

### Probing the internal environment of chromocenters

To gain further insights into the nature of HP1 foci at chromocenters, we compared the fluorescence images resulting from live-cell treatment with **CP-SiR** (**Figure 5A**) and **CP-PA** after photoactivation (**Figure 5B**). In a direct comparison, the fluorescence emission from **CP-PA** was generally fainter due to non-quantitative photoconversion. The intensity-normalized confocal images between the two probes were however very similar, indicating a comparable distribution within the cell and preferential accumulation at HP1-dense foci.

**Figure 5:**
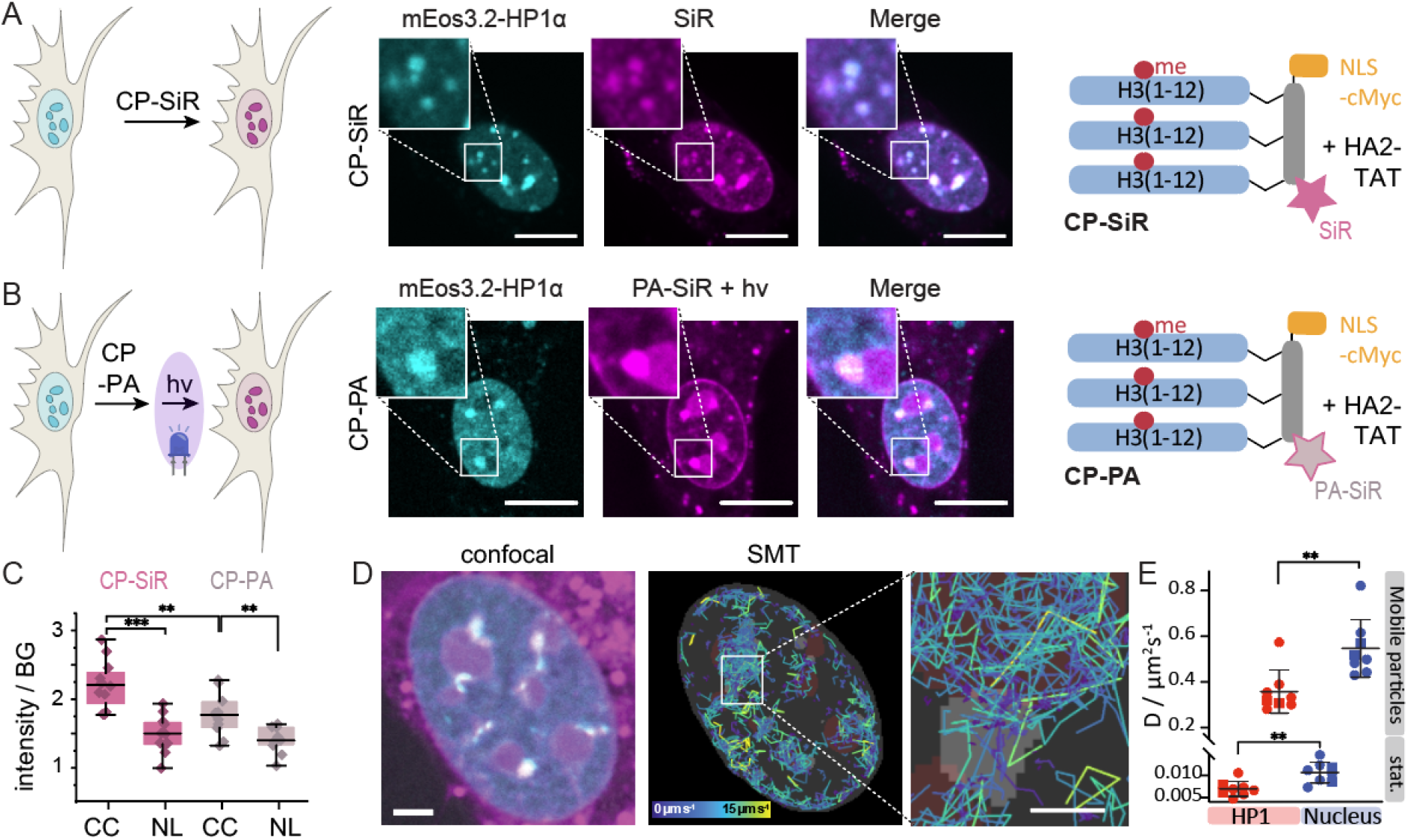
A photoactivatable probe to investigate chromocenter properties. **A**) Heterochromatin staining using **CP-SiR. B**) Heterochromatin staining using **CP-PA** after photoactivation. PA-SiR after 405 nm photoactivation (magenta), HP1α-mEOs3.2 fluorescence (cyan). **C**) **CP-SiR** compared to **CP-PA** fluorescence intensity after photoactivation in chromocenters (CC), compared to nucleoli (NL). **D**) Single-molecule tracks inside the nucleus, colored by velocity. HP1α foci from the reference channel are shown in grey, approximate positions of nucleoli are depicted in dark red. Fast-diffusing molecules are mostly found outside HP1α foci. Molecules that get immobilized after photoactivation have stationary tracks, colored in purple. **E**) Diffusion coefficient estimates for molecules within HP1α foci and in the rest of the nucleus are based on analyzing jump distances in live cells (N=8, 10000 frames). Tracks recorded in different imaging wells are distinguished by the circle and square symbols. Error bars represent standard deviation. Scale bars: A,B = 10 μm, D = 3 μm, inset: 1 μm.

For a deeper analysis, we compared sub-nuclear fluorescence from **CP-SiR**, which indicates peptide probe distribution, to fluorescence emission from **CP-PA**, the photoactivation of which depends on local polarity. In particular, we compared **CP-SiR** and **CP-PA** fluorescence intensity in chromocenters to nucleoli. The nucleolus is understood to be a dense, multi-layered, biomolecular condensate formed by LLPS of proteins including fibrillarin and nucleophosmin, together with RNA^42^, and should thus be a suitable reference point for a phase-separated compartment. Background-normalized emission from **CP-SiR** was used to gauge the degree of accumulation of the peptide probe in chromocenters or nucleoli. We then compared these values to **CP-PA** emission after photoactivation (**Figure 5C**). This analysis revealed that nucleoli showed similar brightness between the two probe types, but chromocenters exhibited a lower degree of fluorescence arising from **CP-PA** than **CP-SiR** (**Figure 5C, Figure S9**). This observation suggests that **CP-PA** is converted less efficiently to a fluorescent state in chromocenters than in phase-separated compartments such as nucleoli, which present good conditions for **CP-PA** photoconversion to a fluorescent state (**Figure 4C-G**). The ∼30% lower fluorescence conversion of **CP-PA** in chromocenters indicates that this environment is comparatively more polar to nucleoli, arguably because of lower protein density.

To determine the diffusional properties of the **CP-PA** probe, we further photoactivated a small subset of molecules and tracked them individually. We grouped the single-molecule tracks according to whether they occurred primarily within or outside chromocenters (**Figure 5D, Figure S10**). Both inside and outside of chromocenters, we could observe mobile, fast diffusing molecules and molecules diffusing at a much slower speed, the latter most likely arising from crosslinking to nearby proteins. Displacements in tracks were fitted assuming two diffusing particle species (high- and low-mobility, **Figure S11**), resulting in approximate diffusion coefficients. This result allowed us to compare the mobility of the probe inside and outside of chromocenters, revealing significantly larger probe mobility outside chromocenters for both high- and low-mobility molecules (**Figure 5E**). Compared to SMT experiments *in vitro*, even the low-mobility molecules showed a ∼20-fold higher diffusion coefficient. This result demonstrates that *in vitro* generated pHP1α condensates do not recapitulate key features of chromocenters and other heterochromatin foci in 3T3 cells, as the latter are more polar (or less dense) and more dynamic overall. Moreover, the fact that the fraction of mobile molecules is higher in chromocenters than in pHP1α condensates generated *in vitro* suggests that more CP-PA molecules are trapped by water as a nucleophile, as opposed to pHP1α. This observation is consistent with our conclusion that chromocenters are less protein-dense than pure pHP1α droplets.

Insight into the organization of interphase chromatin into functional, spatially delimited domains is important for an understanding of gene regulation. Here, we developed peptide-based, cell-permeable probes that can be targeted to a distinct chromatin state by PTM-dependent interaction with reader domains. As the probes are modular, we expect that in future studies, we can target them to other chromatin states, using differently modified histone peptides as targeting sequences. Together, we believe that these probes represent useful tools to image defined chromatin states or to target further chemical functionalities to chromatin sub-compartments.

The nature of heterochromatin organization is of great interest, and multiple models have been put forward. HP1 proteins bind chromatin and, through cross-bridging of nucleosomes, can form dense compartments of compact chromatin^15,16,23^. On the other hand, HP1 proteins efficiently phase-separate *in vitro* via phosphorylation-driven interactions^12^, locally alter chromatin structure^43^, and can stably organize DNA into phase-separated domains^44^. Moreover, such phase-separation behavior has been identified as a driving force for heterochromatin establishment in Drosophila embryos^19^. Maturation of heterochromatin foci coincides with a loss of pure liquid behavior and an increase in stably associated HP1 proteins, resulting in regions of mixed character, containing stably chromatin-bound HP1 and liquid-like domains in specific cell states^19^. We employed the polarity sensitive dye PA-SiR to investigate the internal environment of chromocenters. In 3T3 fibroblasts, **CP-PA** exhibited a similar distribution than the **CP-SiR** probe. However, the probe was less efficiently photoactivated in chromocenters than expected from its behavior in phase-separated pHP1α droplets in *in vitro* studies, indicating a different physicochemical environment. Importantly, the PA-SiR dye allowed for super-resolution imaging and SMT. Dynamic experiments showed that **CP-PA** diffusion was increased in cells compared to *in vitro* pHP1α droplets, providing further evidence that the mature heterochromatin in chromocenters in 3T3 fibroblasts are less apolar and are more heterogeneous than the *in vitro* droplets. Together, these observation supports a model where pHP1α phase separation is not the sole organizing principle in mature cells. Nevertheless, LLPS may contribute to heterochromatin organization in combination with a more gel-like cross-bridged assembly of modified chromatin, HP1 molecules and other heterochromatin proteins.

In summary, our studies add a new tool for the investigation of sub-cellular chromatin compartments, providing functionality to target chromatin compartments using multivalent, modified peptides, image the chromatin states, and, via the application of environmentally sensitive dyes, probe their internal organization.

## Methods

### Automated Solid Phase Peptide Synthesis (SPPS)

General protocol: All peptides were synthesized on a 0.1 mmol scale on a the Tribute peptide synthesizer (PTI) using standard FMOC-chemistry, using Fmoc-aa-hydrazine-Cl-Trt-resin or Rink amide resin (see **Supplementary text** for details). Purification of the crude peptide was achieved with preparative RP-HPLC. Fractions were analyzed, pooled and lyophilized. Purified peptides ware analyzed and characterized by analytical RP-HPLC and ESI-MS (see **Figures S1 – S6**).

### One-pot SiR-Mal or PA-SiR-Mal cysteine labeling and CuAAC reactions (Figure 2)

Scaffold peptide (1 eq., 200 nmol) was dissolved in 50 mM phosphate buffer pH 7.3 to a final concentration of 0.5 mM. A 20 mM SiR-mal solution in DMSO (1.2 eq., 240 nmol) was added, the reaction was agitated at room temperature for 15 min. Alkyne-containing histone tail peptide was added (2 equiv.), followed by addition of a 100 mM solution of CuSO_4_ (4 eq., 800 nmol) and a 500 mM solution of sodium ascorbate (20 eq., 4 μmol). The reaction mixture was agitated at room temperature for 5 min and immediately purified by semi-preparative RP-HPLC. Product identity was confirmed by RP-HPLC and ESI-MS analysis (**Figures 2, S4**).

### Cell culture and imaging

NIH 3T3 cells (ATCC CRL-1658) were seeded in 6-well plates to a density of 2×10^5^ cells/well and left to attach overnight at 37°C in 1.5 ml DMEM-Glutamax supplemented with 10% FBS (Gibco) and penicillin-streptomycin (10 <u>μg</u>/ml, Gibco). The following day, medium was replaced with 1.5 ml DMEM-Glutamax supplemented with 10% FBS (Gibco). Lipofectamine 2000 (Invitrogen) was used for transfection following manufacturer’s instructions. Transfection reaction used 12 μg of pcDNA3- mEOS3.2-HP1α plasmid and a ratio of 1:4 μg DNA to Lipofectamine 2000. After 4 h incubation, the medium was replaced and left overnight at 37°C. The following day, cells were detached using Trypsin-EDTA and counted before being seeded into glass bottom 96-well plates (Mattek) at density of 5×10^3^ cells/well in 100 μl DMEM-Glutamax supplemented with 10% FBS (Gibco) and penicillin-streptomycin (10 μg/ml, Gibco). Cells were left to attach overnight at 37°C. Cells were washed with 100 μl warm PBS (Gibco) and incubated with probe (5 μM) while being co-incubated with endosolytic peptide^4^ to help cellular uptake, after cells were washed with incubated with DMSO diluted probe in Optimem medium (Gibco).

Images (pHP1α droplets or live cells) were acquired on a confocal microscope LSM 510 Meta (Zeiss), using a C-Apochromat 63x/1.2 water corrected objective. Alternatively, we used a Nikon Ti2 eclipse microscope equipped with a Mad City Labs piezo motorized stage and Okolab incubator and sample holders. See **Supplementary text** and **Figure S10** for a detailed confocal and single-molecule imaging protocol.

### Purification of pHP1α

Phosphorylated HP1α was expressed as described previously^3^. BL21(DE3) E. coli cells were transformed with a pRSFDuet vector containing his6-tagged human HP1α and mouse casein kinase 2 (CK2). The cells were lysed by sonication and purified over via immobilized metal affinity chromatography (IMAC) over a Ni:NTA resin as per manufacturer’s instructions. Subsequently, the his6-tag was removed by thrombin cleavage, and pHP1α was further purified via size-exclusion chromatography (SEC) using a S200 column in SEC buffer (50 mM HEPES pH 7.3, 150mM NaCl, 1mM DTT). The eluted pure fractions were combined, concentrated, glycerol was added to 30% and the protein was stored, snap-frozen as 10 μL aliquots at – 80°C.

### *In vitro* pHP1α phase separation and confocal microscopy

Working stock solutions of peptide conjugates **CP-SiR, CP’-Sir, CP-PA or CP’-PA** were made by dilution with H_2_O at 400 nM concentration. For the use of PEG, PEG 6000 was dissolved in H_2_O to give a 20 %w PEG stock solution and 60 μM phosphorylated HP1α protein stock solution was also prepared. PEG solution (1 μL) and pHP1α stock solution (2 μL) were mixed in an Eppendorf vial by pipetting, resulting in a solution with phase-separated droplets. After 2 minutes, peptide conjugate (1 μL) was added to the mixture, was homogenized by further pipetting, and the total sample (4 uL) was placed on a cover slide for imaging. Time elapsed after mixing for each image was calculated from recorded mixing time and the time of capture. Measurements with similar times after mixing were grouped to triplicates (**Figures 4, S8**). See **Supplementary text** for a detailed imaging protocol.

## Acknowledgments

We thank Prof. Stefan Matile for the diselenolane-NHS ester. B. F. thanks the NCCR Chemical Biology and EPFL for financial support. P. R.-F thanks the NCCR Chemical Biology, the Swiss National Science Foundation (grant 206021_189631), and the European Research Council (ERC Starting Grant 801572, HDPROBES) for financial support.

## TOC Graphic

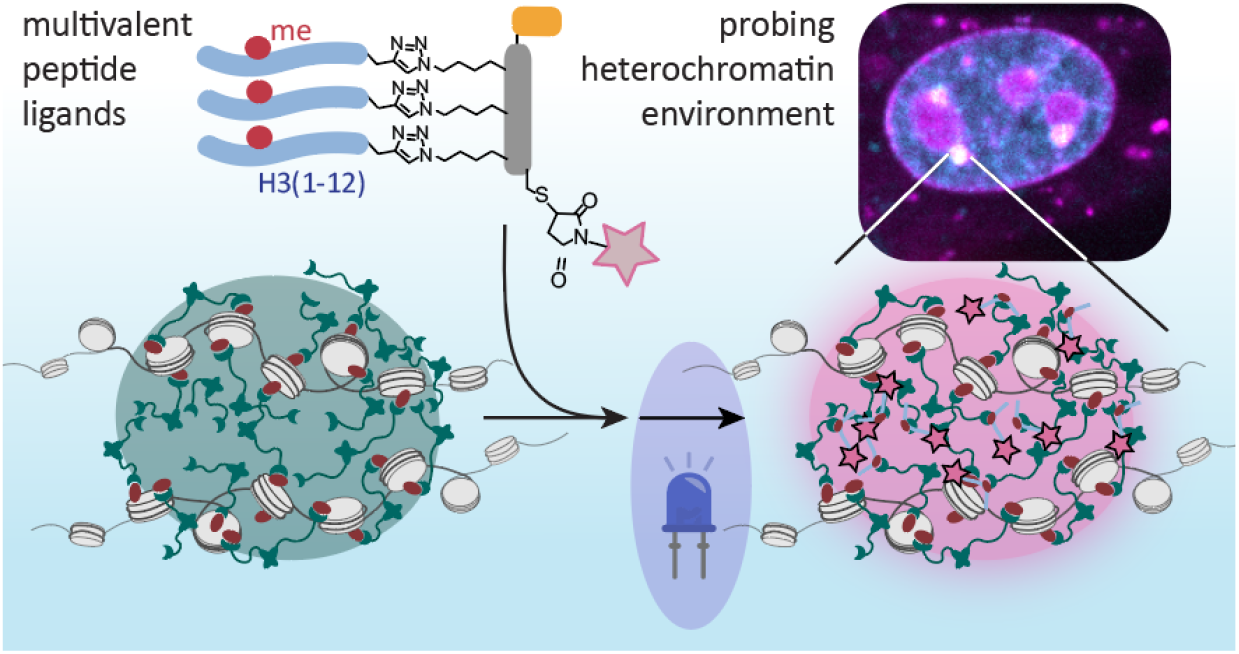

