## Supplementary Materials for "Multivalent peptide ligands to probe the chromocenter microenvironment in living cells"

All solvents and reagents were purchased from commercial sources and used without further purification. Solvents were procured from Sigma-Aldrich, Acros, Fluorochem, and TCI and used as received. Fmoc-L-Lys(Boc,Me)-OH was from Iris Biotech (Marktredwitz, Germany). Fmoc-L-Lys(Me)$_3$-OH chloride, Fmoc-Lys(N$_3$)-OH, Fmoc-Pra-OH and Rink amide MBHA were from GL Biochem (Shanghai, China). Boc-L-thiazolidine-4-carboxylic acid was from Bachem (Bubendorf, Switzerland). All other amino acid derivatives, 2-chlorotrityl chloride resin and 2-(7-Aza-1H-benzotriazole-1-yl)-1,1,3,3-tetramethyluronium hexafluorophosphate (HATU) were purchased from Novabiochem, Merck (Darmstadt, Germany). N,N-Dimethylformamide (DMF), N,N-diisopropylethylamine (DIEA), piperidine and phenol were from Acros Organics (Geel, Belgium). 2-(1H-benzotriazol-1-yl)-1,1,3,3-tetramethyluronium hexafluorophosphate (HBTU) was from Protein Technologies Inc. (Tucson, USA). (6-Chlorobenzotriazol-1-yl)-N,N,N′,N′-tetramethyluronium hexafluorophosphate (HCTU) was from Carl Roth GMbH (Karsruhe, Netherlands). Hydrazine monohydrate was purchased from Alfa Aesar (Heysham, UK), acetonitrile (ACN) from Avantor Performance Materials (USA). 4-mercaptophenyl acetic acid (MPAA), Diethylether, acetic anhydride, phenylsilane, tetrakis(triphenylphosphine)palladium(0), hydroxybenzotriazole (HOBt), 5,5′-Dithiobis(2-nitrobenzoic acid) (DTNB), Dithiothreitol (DTT), silver acetate, trifluoroacetic acid (TFA), dichloromethane (DCM), triisopropylsilane (TIS), ethanedithiol (EDT), thioanisole, sodium nitrite, L-glutathione reduced (GSH), sodium diethyldithiocarbamate trihydrate and methyl thioglycolate (MTG), Tris(2-carboxyethyl)phosphine hydrochloride (TCEP), dimethyl sulfoxide (DMSO), sodium ascorbate, Copper sulfate, were from Sigma Aldrich (Taufkirchen, Germany). All other commonly used chemical reagents and buffer components were from Applichem (Darmstadt, Germany) and Fisher Scientific (Reinach, Switzerland). Restriction enzymes, Q5 DNA polymerase, Phusion polymerase, T5 exonuclease, Taq DNA ligase, dTNPs, DNA ladders and DNA loading dyes were from New England Biolabs (Ipswich, MA, USA). QiaQuick spin column for PCR purification, gel extraction and nucleotide removal as well as QiaPrep spin columns for miniprep plasmid purification were from Qiagen (Hilden, Germany). Primers were ordered from and synthesized by Integrated DNA technologies (Leuwen, Belgium) and gene sequencing was performed at GATC Biotech (Constance, Germany). Chemicals and solutions for preparation of agarose and SDS polyacrylamide gels (agarose, acrylamide, Precision Plus Protein™ All Blue Prestained Protein Standard) were purchased from BioRad (Hercules, CA, USA). Mammalian cell culture media and components, such as Dulbecco's modified eagle's medium (DMEM)-GlutaMAX, Opti-MEM reduced serum medium, Phosphate-buffered saline (PBS), Trypsine-EDTA, Penicillin-Streptomycin (Pen-Strep) and Lipofectamine 2000 were from Gibco-Invitrogen (Basel, Switzerland).

### Organic synthesis methods – Synthesis of the photoactivatable probe

**General methods.** NMR spectra were recorded on Bruker 400-600 MHz instruments. $^{1}$H NMR chemical shifts are reported in ppm relative to SiMe$_4$ (δ = 0) and were referenced with respect to residual protons of the solvent (δ = 7.26 for CDCl$_3$, and δ = 2.50 for dimethyl sulfoxide, δ = 1.94 for acetonitrile). Coupling constants are reported in Hz. $^{13}$C NMR chemical shifts are reported in ppm relative to SiMe$_4$ (δ = 0) and were referenced internally with respect to solvent signal (δ = 77.16 for CDCl$_3$ and δ = 39.52 dimethyl sulfoxide, δ = 118.26 for acetonitrile). Preliminary peak assignments are based on calculated chemical shift, multiplicity, and HSQC and HMBC spectra. Liquid chromatography coupled mass spectrometric (LCMS) analysis was done using Shimadzu LCMS-2020 spectrometer by using electrospray ionization (ESI). IUPAC names of all compounds were generated using PerkinElmer ChemDraw 20.1. Preparative reverse phase purification was done using Agilent 1260 Infinity HPLC with Agilent Zobrax 300SB-C18 5 µM column (9.4 × 250 mm).

Absorbance measurements were made using Thermo Scientific Multiskan Sky spectrophotometer with Thermo Scientific µdrop insert (N12391). In all cases, miliQ $H_2O$ and optical grade DMSO were used to prepare stock solutions, dilutions. All stock solutions were stored at -15 °C. Before use, vials were kept on ice and were used immediately after thawing. 5′-alkynyl-silicon rhodamine diazoindanone **PA-SiR.1** was synthesized according to previously reported procedures.[1] Synthesis of azidopropionic acid was performed according to previous protocols. [2]

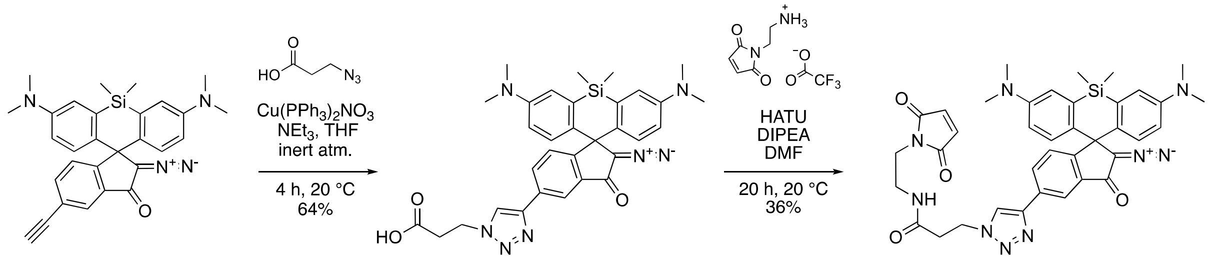


**PA-SiR.1** **PA-SiR.2** **PA-SiR-Mal**

**Supporting Scheme 1.** Synthesis of photoactivatable dye with maleimide functionality.

**3-(4-(2'-Diazo-3,7-bis(dimethylamino)-5,5-dimethyl-3'-oxo-2',3'-dihydro-5*H*-spiro[dibenzo[*b*,*e*]siline-10,1'-inden]-5'-yl)-1*H*-1,2,3-triazol-1-yl)propanoic acid (PA-SiR.2)**

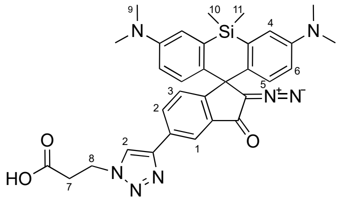

5′-alkynyl-silicon rhodamine diazoindanone (12 mg, 25.2 µmol, 1 eq.) was placed in a flask with azido propionic acid (4.35 mg, 37.8 µmol, 1.5 eq.). The flask was sealed with a septum, then it was flushed with $N_2$ and starting materials were dissolved in tetrahydrofuran (6 mL). The solution was purged by bubbling $N_2$ through the solution. Catalyst $Cu(PPh_3)_2NO_3$ (4 mg, 6.15 µmol, 0.244 eq.) and base triethylamine (1.75 µL, 12.6 µmol, 0.5 eq.) were added in tetrahydrofuran (6 mL). The solution was stirred for 4 h at 20 °C, when full conversion was observed. Product was isolated by normal phase column chromatography (0–10% MeOH, and 0–10% EtOAc in $CH_2Cl_2$, MeOH:EtOAc 1:1, 3 g $SiO_2$), 9.5 mg (64%). $^1$H NMR (500 MHz, $CD_3CN$) δ 8.04 (s, 1H), 7.85 – 7.77 (m, 2H), 7.36 (s, 1H), 6.98 (d, *J* = 3.0 Hz, 2H), 6.89 (d, *J* = 9.0 Hz, 2H), 6.68 (dd, *J* = 9.0, 3.0 Hz, 2H), 4.53 (t, *J* = 6.6 Hz, 2H), 2.91 (s, 12H), 2.88 (t, *J* = 6.6 Hz, 2H), 0.68 (s, 3H), 0.50 (s, 3H) ppm. $^{13}$C NMR (126 MHz, $CD_3CN$) δ 187.94, 172.13, 159.73, 149.84, 146.46, 138.08, 135.57, 134.56, 133.48, 130.45, 126.12, 124.03, 123.19, 122.15, 116.91, 116.23, 77.94, 58.09, 46.59, 40.44, 34.45, 0.70, 0.26 ppm.

**3-(4-(2'-Diazo-3,7-bis(dimethylamino)-5,5-dimethyl-3'-oxo-2',3'-dihydro-5H-spiro[dibenzo[b,e]siline-10,1'-inden]-5'-yl)-1H-1,2,3-triazol-1-yl)-N-(2-(2,5-dioxo-2,5-dihydro-1H-pyrrol-1-yl)ethyl)propanamide (PA-SiR-Mal)**

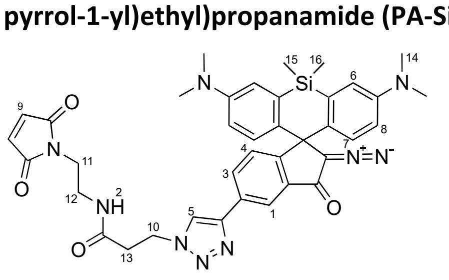

NN-SiR carboxylic acid **xx** (9.17 mg, 15.5 µmol, 1 eq.) was dissolved in dry N,N-dimethylformamide (6 mL) in a septum sealed flask. HATU (9 mg, 23.7 µmol, 1.53 eq.) and DIPEA (15.4 µL, 93 µmol, 6 eq.) were added, and the resulting mixture was stirred at rt for 20 min (formation of the activated ester monitored by TLC). Afterwards, the first portion (4 mg) of N-(2-aminoethyl)maleimide, trifluoroacetate salt (4+2 mg, 23.6 µmol, 1.52 eq.) was added to the reaction, and the solution was stirred at rt for 4 h. To reach full conversion, a second portion of amine reagent (2 mg) was added. The reaction was diluted with DCM and washed with brine (3×). The aqueous layer was re-extracted with DCM (1×) and the combined organic layers were dried over sodium sulfate and concentrated under reduced pressure. Drying on high vacuum was necessary to evaporate residual DMF. The product was purified by normal phase column chromatography (0–10% MeOH, and 0–10% EtOAc in $CH_2Cl_2$, MeOH:EtOAc 1:1, 2 g $SiO_2$) then further purified with reverse phase chromatography (5–40% MeCN in $H_2O$, over 40 min, C18 column) to yield 4 mg (36%) pure compound.

$^1$H NMR (400 MHz, DMSO) δ 8.53 (s, 1H), 8.06 (t, *J* = 6.1 Hz, 1H), 7.91 (dd, *J* = 8.0, 1.4 Hz, 1H), 7.84 (d, *J* = 8.0 Hz, 1H), 7.37 (t, *J* = 0.9 Hz, 1H), 6.94 (d, *J* = 2.8 Hz, 2H), 6.89 (s, 2H), 6.75 (d, *J* = 9.0 Hz, 2H), 6.70 (dd, *J* = 9.1, 2.8 Hz, 2H), 4.49 (t, *J* = 6.9 Hz, 2H), 3.39 (t, *J* = 5.8 Hz, 2H), 3.14 (q, *J* = 6.0 Hz, 2H), 2.89 (s, 12H), 2.60 (t, *J* = 6.9 Hz, 2H), 0.66 (s, 3H), 0.47 (s, 3H) ppm. $^{13}$C NMR (151 MHz, DMSO) δ 186.97, 171.44, 169.56, 158.82, 148.83, 145.25, 137.50, 134.86, 134.58, 133.61, 132.54, 129.59, 125.69, 123.61, 123.48, 121.41, 116.22, 115.82, 77.14, 57.16, 46.31, 39.95, 37.44, 37.38, 35.65, 31.18, 1.35, 0.46 ppm. HRMS (ESI/QTOF) m/z: $[M + Na]^+$ Calcd. for $C_{38}H_{39}N_9NaO_4Si^+$ 736.2786; Found: 736.2787.

### Peptide synthesis

#### Automated Solid Phase Peptide Synthesis (SPPS)

General protocol: All peptides were synthesized by the Tribute peptide synthesizer (PTI) on the previously prepared Fmoc-aa-hydrazine-Cl-Trt-resin or Rink amide resin to yield peptides with C-terminal hydrazide or amide, respectively. The syntheses were performed on 0.1 mmol scale using Fmoc chemistry. The standard base-resistant groups employed to protect amino acid side chains are listed below: Arg(Pbf), Lys(Boc), Thr(tBu), Gln(Trt), Ans(Trt), Asp(OtBu), His(Trt), Cys(Trt), Ser(tBu), Tyr(tBu), Glu(tBu). In addition, the following orthogonal protecting groups were employed: Cys(Acm), Lys(Alloc), Lys(Me), Lys(Me3), Lys(N3), Gly(Pra), Glu(OAll). To maximize synthesis yield, amino acids were double coupled and pseudoproline dipeptide building blocks were used where necessary.

Briefly, the N-terminal Fmoc-group was deprotected by incubating the resin with 20% (v/v) piperidine in DMF. Activation of amino acid (0.5 mmol, 5 eq.) was achieved by addition of HBTU or HCTU (0.48 mmol, 4.8 eq.) and DIEA (1 mmol, 10 eq.). The coupling step was performed by adding the activated amino acid to the resin, followed by 30 min incubation at room temperature. When the full-length peptide was assembled, the peptidyl-resin was washed with DMF, DCM and MeOH and dried under vacuum.

Peptides were cleaved from the resin using either 95% TFA, 2.5% TIS, 2.5% $H_2O$ or 87.5% TFA, 5% phenol, 5% thioanisole, 2.5% ethanedithiol, 5% $H_2O$. The crude peptide was precipitated by addition of ice-cold diethyl ether, recovered by centrifugation, dissolved in 50% (v/v) acetonitrile in $H_2O$, flash-frozen and lyophilized.

#### Synthesis of cR10

The linear peptide was synthesized by automated Fmoc-SPPS as reported above, on rink amide resin, with alternating Fmoc-L-Arg(Pbf)-OH and Fmoc-R-Arg(Pbf)-OH amino acids. Alloc/Allyl deprotection: the peptidyl-resin was swollen for 30 min in DCM. 3 mL of dry DCM and $PhSiH_3$ (5 mmol, 50 eq.), followed by $Pd(PPh_3)_4$ (0.05 mmol, 0.5 eq.) in 0.5 mL dry DCM were added to the resin and incubated for 30 min at room temperature. The resin was washed with DCM and the deprotection reaction with $PhSiH_3$ and $Pd(PPh_3)_4$ was repeated two more times. The resin was thoroughly washed with DCM followed by washing with 0.5% (v/v) DIEA in DMF; 0.5% (w/v) sodium-diethyldithiocarbamate in DMF; 50% (v/v) DCM in DMF; 0.5% (w/v) HOBt in DMF and intensively washed with DMF. On-resin cyclization: glutamate side chain carboxylic acid was activated with a 0.5 M stock solution of HATU in DMF (1 eq., 0.1 mmol) and DIEA (8 eq., 0.8 mmol). The cyclization reaction was allowed to occur for 2h, under $N_2$ bubbling at room temperature. After extensive DMF washes, 2x500 μL 20% (v/v) piperidine were added to the resin for 5 min at room temperature to remove the N-terminal Fmoc. The resin was extensively washed with DMF, DCM and MeOH and dried under vacuum. The peptide was cleaved and purified as described above (**Figure S5A**).

#### Heterodisulfide formation (Figures S3, S5)

Cys-Acm deprotection: In a typical reaction, peptide probe (e.g. **$(T2)_3$-SN3-SiR**) was dissolved in 50% AcOH to a concentration of 100 mM. AgOAc was dissolved in 50% AcOH to a concentration of 30 mM, and mixed to the peptide solution in a 1:1 ration (final concentrations: peptide 0.5 mM, AgOAc 15 mM). The solution was subsequently incubated at 37°C for 3h under agitation. The reaction progress was monitored by HPLC analysis. DTNB activation: To the reaction mixture, containing the deprotected peptide, an equal volume of 60 mM 5,5′-Dithiobis(2-nitrobenzoic acid) (DTNB) in 6M GdmHCl, 0.2 M phosphate buffer, pH 7.4, was added, followed by vigorous mixing. The reaction mixture was incubated for 5 min, followed by centrifugation. The supernatant, containing the TNB-modified peptide, was removed and diluted 5-fold with buffer pH 3, containing 5 M GdmHCl and purified over semipreparative HPLC. Heterodisulfide-formation: The purified TNB-modified peptide was dissolved in 6M GdmHCl, 0.2 M phosphate buffer, pH 6 to a concentration of about 0.5 mM. Cysteine-containing peptides (e.g. cR10-SH, **Figure S5**) was dissolved in the same buffer and added in 3 eq. excess, followed by rapid mixing. After 2 min incubation, the reaction mixture was 10-fold diluted with the same buffer, and immediately purified by semipreparative HPLC. Pure peptide fractions were analyzed by analytical HPLC, ESI-MS, lyophilized and stored as dried powder at – 20°C.

**SeL labeling (Figure S6)**
5 μL of 100 mM SeL-NHS solution in DMSO (2 eq., 500 nmol) were mixed with 2.5 μL of 100 mM *N*-(2-aminoethyl)maleimide in DMSO (1 eq., 250 nmol). The solution was diluted to 25 μL with 100 mM phosphate buffer pH 8.2. After 1h incubation at room temperature, excess glycine (5 eq., 1.25 μmol) was added to quench the remaining SeL-NHS. Acm-deprotected peptide probe (1 eq., 100 nmol) was dissolved in 100 mM phosphate buffer, pH 8.2, to a final concentration of 0.5 mM in a siliconized 1.5 mL tube. Crude SeL-mal (2 eq., 200 nmol) was added and the reaction mixture was vortexed and shaken at room temperature for 15min. The SeL-probe peptide was purified by semipreparative RP-HPLC and lyophilized. Product identity was confirmed by RP-HPLC and ESI-MS analysis.

### *In vitro* phase-separation experiments and imaging

**Imaging methods**
Images (pHP1α droplets or live cells) were acquired on a confocal microscope LSM 510 Meta (Zeiss), using a C-Apochromat 63x/1.2 water corrected objective. Imaging was performed using the following parameters: pixel time: 1.28 μs, excitation lasers: 488 nm (13% emission power) and 633 nm (25% emission power), detector gain: 550.
Alternatively, we used a Nikon Ti2 eclipse microscope equipped with a Mad City Labs piezo motorized stage and Okolab incubator and sample holders. For confocal imaging, an Omicron Laserhub served as light source, connected to Yokogawa W1 spinning disc confocal scanning unit. The objective used was a Nikon 1.49 NA 100x TIRF Apo Plan SR objective, and fluorescence signal was recorded by Telendyne Prime 95B sCMOS cameras. For single-molecule imaging Omicron Laserhub light source was used with a Nikon N-STORM TIRF illuminator and a Nikon 1.49 NA 100x TIRF Apo Plan SR objective. The image was projected onto an Andor iXon 888 ultra EMCCD camera. All lasers and camera shutters were controlled by National Instruments NIDAQ controller unit. All microscope laser powers reported correspond to intensities at the end of the optical fiber, before entering the microscope unit. Excitation laser power at sample plane were measured for confocal imaging (405 nm: 65 mW at fiber tip = 10.0 mW at sample, 638 nm: 100 mW at fiber tip = 13.1 mW at sample).

**Concentration measurements**
Concentrations of all peptide conjugates containing the SiR dye were determined by UV-Vis and using the published SiR extinction coefficient[1] for concentration calculation. The concentration of peptides **CP-PA** and **CP'-PA** and the photoactivatable maleimide dye **PA-Sir-Mal** were also determined by UV-spectroscopy. CP. The absorption spectrum between 300-400 nm matched for all three compounds due to the absorption of diazoindanone chromophore at this wavelength[2]. The concentration of maleimide dye **PA-Sir-Mal** was measured as 3.8 mM using quantitative NMR, with 1.19 mM acetone used as an internal standard (**Figure S7**).

To measure the concentration of peptide conjugates, a dilution series in DMSO were made from the maleimide dye as reference. Absorption of the peptide conjugates and the maleimide dye precursor matched had identical peaks in the 300-400 nm region due to the diazoindanone chromophore. An absorbance calibration linear was recorded based on absorbance at 319 nm. The concentration of peptide conjugates was determined as **CP-PA** 333 µM and **CP'-PA** 288 µM.

#### *In vitro* pHP1α phase separation and confocal microscopy

Detailed imaging protocol: Before recording each image, the focus was set to 0.8 µm above the plane of glass. This was important because the dye showed significant turn-on at the glass-sample interface. By setting the focal plane 0.8 µm above glass, the signal of the interface was defocused and minimized, while being close enough to the surface to image stationary sedimented droplets.

For imaging of the PA-SiR dye, the following imaging sequence was used:

1. Transmission image in green (50 ms)
2. Short photobleaching using epifluorescent illumination (638 nm, 3 s, 110 mW)
3. Before photoactivation confocal image in far-red (638 nm, 800 ms, 100 mW)
4. Photoactivation of sample by confocal exposure (405 nm, 200 ms, 65 mW)
5. Before photoactivation image in far-red using confocal (638 nm, 800 ms, 100 mW)

The photoactivation with the exposure listed above activated approximately 20-30 % of the total precursor.

Images were segmented on the multichannel image using Ilastik[6] (https://www.ilastik.org/). The segmented particles were analyzed in ImageJ to provide particle sizes. Intensity values were measured on the original images before and after photoactivation, in and outside droplets based on the segmentation by Ilastik. Characterization of the resulting droplets are presented in **Figure S8 E-G.** Control probe **CP'-SiR** showed very similar intensities inside and outside the droplets, leading to non-accurate partial sizes measured, especially in the earlier timepoints during the experiment.
In single molecules acquisitions **Figure S8** acquired 45 min - 2 h after mixing

In experiment **Figure 4 G,E** the full sample was illuminated with Roithner LED (405 nm, 150 s, 2 mW $cm^{-2}$) to perform exhaustive photoactivation of all the precursor.

### Cell culture and imaging protocols

#### Cell culture imaging protocol and data analysis

For *confocal imaging* the following confocal image sequence was recorded:

1. Transmission image in green (50 ms)
2. Before photoactivation confocal image in far-red (638 nm, 300 ms, 100 mW)
3. Reference image by confocal exposure (488 nm, 200 ms, 24 mW)
4. Photoactivation of sample by confocal exposure (405 nm, 200 ms, 65 mW)
5. Before photoactivation image in far-red using confocal (638 nm, 300 ms, 100 mW)
   In some filed of views photoactivating image (4) in the sequence was omitted.

Fiji software was used to analyze images (see e.g. **Figure 3, S9**). Colocalization analysis of mEos3.2-HP1α and probe-SiR signals was performed on single cells. First, a nuclear mask was obtained by smoothing the mEos3.2-HP1α channel signal with a Median filter and applying the IsoData thresholding algorithm, followed by "Fill holes" command. The nuclear mask was applied to SiR-probe channel to exclude any SiR signal outside of the nucleus from the analysis. Background correction was performed in the mEos3.2-HP1α and probe-SiR channels setting a rolling ball radius of 20 and 50 pixels, respectively. Pearson's correlation coefficients (**Figure 3J**) were calculated with the BIOP JACOBS plugin, using the nuclear mask as a ROI, to exclude pixels outside of the nucleus to be considered in the analysis.

**Single-molecule imaging**

In cells with sufficient nuclear signal in far-red channel and alongside successful transfection detected, single molecule acquisition was recorded (total 8 field of views from 2 wells). Cells were photobleached while the HILO illumination was adjusted (638 nm, approx. 3 min, 110 mW). Then cells were imaged with the illumination sequence displayed in **Figure S10**. Initial reference image was recorded (488 nm, 20 ms, 10-20 mW) while photoactivation of single molecules took place (488 nm, 1 ms, 1 mW). The reference- and activation frame was followed by 100 readout frames to record single molecule positions (Confocal imaging: 638 nm, 300 ms, 100 mW, SMLM: 638 nm, 50 ms, 100 mW). This cycle of activation followed by readout frames was repeated 100 times, resulting in total 10000 frames. The total duration was over 9 minutes, with use of a constant framerate 18.35 frames per second.

To process illumination sequence data, first reference images were denoised with the Pure denoise plugin in ImageJ[5]. Then reference images were segmented using Ilastik[6], trained for each field of view individually. From the resulting segmentations, index images were generated that indicate positions HP1α speckles and the nucleus. Index images were interpolated for all frames with single molecule signal. In the case of few fields of views approximate nucleoli positions could be drawn manually as well. Nucleoli positions were based on confocal images taken before and after single molecule imagining.

Single molecule data (**Figure S10**) was analyzed using TrackMate v6.0.2 with TrackMate extras 0.0.4 addon[7]. Single molecules were detected with difference of Gaussians detector with median filter, then localization outside the nucleus were omitted based on index image. Kept localizations were joined to tracks based on a linear assignment problem algorithm[8] with 1.2 µm linking threshold. Linking thereshold was based on the largest observed single molecule displacements. The probability of false linking assuming random distributions of the median 3 emitters in the nucleus (160 $\mu m^2$) was approximately 5.5%. Gap closing of 1 frame was allowed in case of molecules with similar brightness. Tracks with more than two gaps closing were omitted, and tracks with less than 4 localizations were also removed. Remaining tracks were divided to two subsets: 1, tracks with particles in HP1α speckles and 2, tracks in the nucleus with particles outside HP1α speckles. These subsets of tracks were subjected to diffusion coefficients analysis with Spot-On[9] (**Figure S11**). Displacements histograms were generated for tracks inside HP1 foci (n > 130 per cell) and tracks outside HP1 foci (n > 800 per cell) for each cell separately. Displacement histograms were made for 7 consecutive frames which were fitted assuming 2 diffusing species, resulting in 2 diffusion coefficient values for both inside and outside HP1 foci. For diffusion coefficient fitting XY uncertainty of detections was assumed as 20 nm, based on Gaussian fitting parameters. The depth of optical detection (z) of single molecules was approximated as 0.7 µm.

### Supplementary tables

| Name | Description | Sequence | Mass (Da) |
|---|---|---|---|
| **Targeting peptides** | | | |
| T1 | H3(1-15)K9me3 | ARTKQTARK(me3)STGGKAGG(Pra)-NH2 | 1755.0 |
| T1' | H3(1-15)K9A | ARTKQTARASTGGKAGG(Pra)-NH2 | 1654.8 |
| T2 | H3(1-12)K9me3 | ARTKQTARK(me3)STGSG(Pra)-NH2 | 1528.8 |
| T2' | H3(1-12)K9A | ARTKQTARASTGSG(Pra)-NH2 | 1428.6 |
| **Scaffold peptides** | | | |
| S | scaffold | RGSK($N_3$)GSGSK($N_3$)GSGSCC-NH2 | 1479.6 |
| SN1 | TAT-CPP scaffold | RKKRRQRRR-GSGSK($N_3$)GSGSK($N_3$)GSGSCC(Acm)-NH2 | 2645.0 |
| SN2 | NLS-SV40 scaffold | PKKKRKV-GSGSK($N_3$)GSGSK($N_3$)GSGSCC(Acm)-NH2 | 2187.5 |
| SN3 | NLS-cMyc scaffold | PAAKRVKLD-GSK($N_3$)GSK($N_3$)GSK($N_3$)GSCC(Acm)-NH2 | 2312.6 |
| **Other peptides** | | | |
| HA2-TAT | endosomolytic peptide | CGLFEAIAEFIENGWEGLIEGWYGGRKKRRQRRR-NH2 | 4095.8 |

**Table 1:** Overview over all peptide components used for probe synthesis.

#### Table 2

| Name | Alt. name | Scaffold | Chromatin targeting | Cell entry | Detection | MW |
|---|---|---|---|---|---|---|
| (T1)$_2$-SN1-SiR | | (**K**GSGS)$_2$-CC(Acm)-NH2 | T1: H3(1-15)K9me3 | N1: TAT-CPP | SiR | 6749.8 |
| (T1)$_2$-S-SiR | | (**K**GSGS)$_2$-CC(Acm)-NH2 | T1: H3(1-15)K9me3 | N/A | SiR | 5584.4 |
| (T1)$_2$-SN2-SiR | | (**K**GSGS)$_2$-CC(Acm)-NH2 | T1: H3(1-15)K9me3 | N2: NLS-SV40 | SiR | 6293.4 |
| (T1)$_3$-SN3-SiR | | (**K**GSGS)$_3$-CC(Acm)-NH2 | T1: H3(1-15)K9me3 | N3: NLS-cMyc | SiR | 8172.5 |
| (T1)$_3$-SN3C2-SiR | | (**K**GSGS)$_3$-CC(Acm)-NH2 | T2: H3(1-12)K9me3 | N3: NLS-cMyc / C2: cR10 | SiR | 8399.8 |
| (T2)$_2$-SN2C3-SiR | | (**K**GSGS)$_2$-CC(Acm)-NH2 | T2: H3(1-12)K9me3 | N2: NLS-SV40 / C3: SeL | SiR | 6644.5 |
| T2-SiR | | N/A | T2: H3(1-12)K9me3 | N/A | SiR | 2082.1 |
| (T2)$_3$-SN3-SiR | CP-SiR | (**K**GSGS)$_3$-CC(Acm)-NH2 | T2: H3(1-12)K9me3 | N3: NLS-cMyc | SiR | 7493.7 |
| (T2')$_3$-SN3-SiR | CP'-SiR | (**K**GSGS)$_3$-CC(Acm)-NH2 | T2': H3(1-12)K9A | N3: NLS-cMyc | SiR | 7193.1 |
| (T2)$_3$-SN3-PA | CP-PA | (**K**GSGS)$_3$-CC(Acm)-NH2 | T2: H3(1-12)K9me3 | N3: NLS-cMyc | PA-SiR | 7613.8 |
| (T2')$_3$-SN3-PA | CP'-PA | (**K**GSGS)$_3$-CC(Acm)-NH2 | T2': H3(1-12)K9A | N3: NLS-cMyc | PA-SiR | 7313.2 |

**Table 2:** Overview over all heterochromatin targeting peptide probes (see also **Figure 3**)

### Supplementary Figures

#### Supplementary Figure 1

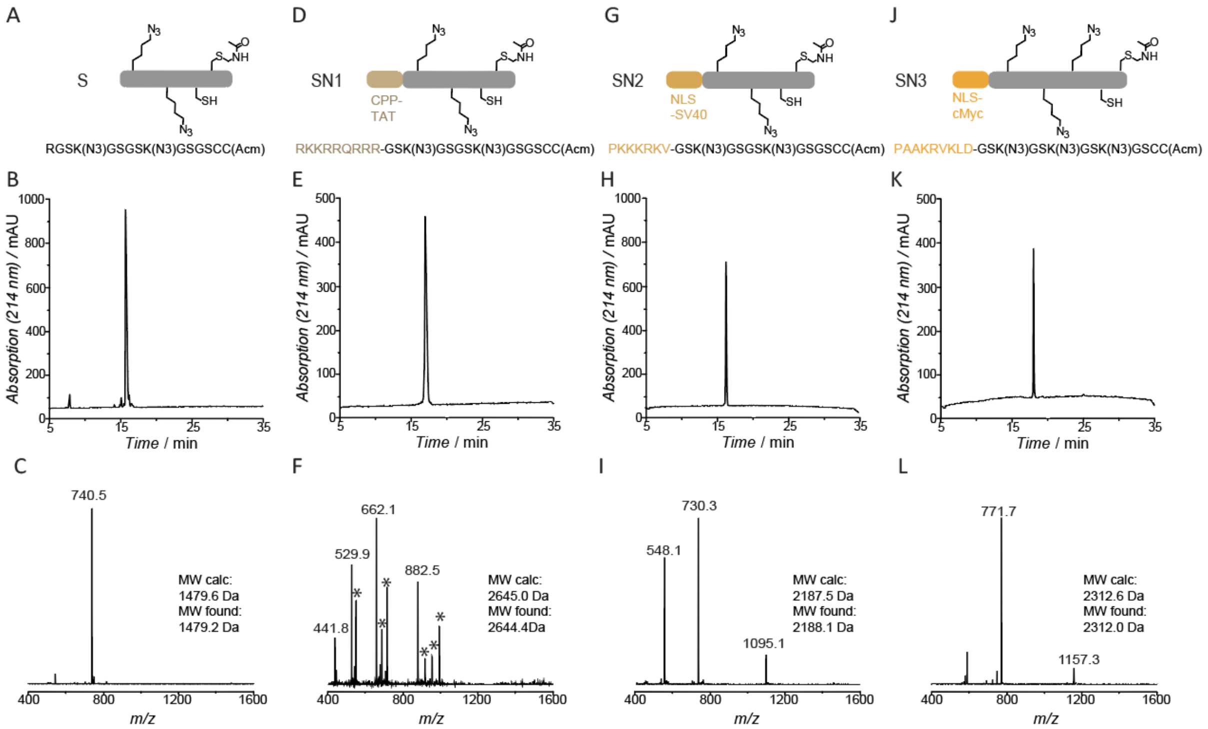


**Figure S1 – Analytics of scaffold peptide building blocks. A**) Sequence of the scaffold peptide **S. E**) Analytical RP-HPLC of scaffold peptide **S**. **F**) ESI-MS analysis of scaffold peptide **S** (MW calculated: 1479.6 Da, MW found: 1479.2 Da). **D**) Sequence of the scaffold peptide **SN1. E**) Analytical RP-HPLC of scaffold peptide **SN1**. **F**) ESI-MS analysis of scaffold peptide **SN1** (MW calculated: 2645.0 Da, MW found: 2644.4 Da). Asterisks are TFA adducts. **G**) Sequence of the scaffold peptide **SN2. H**) Analytical RP-HPLC of scaffold peptide **SN2**. **I**) ESI-MS analysis of scaffold peptide **SN2** (MW calculated: 2187.5 Da, MW found: 2188.1 Da). **J**) Sequence of the scaffold peptide **SN3. K**) Analytical RP-HPLC of scaffold peptide **SN3**. **L**) ESI-MS analysis of scaffold peptide **SN3** (MW calculated: 2312.6 Da, MW found: 2312.0 Da).

### Supplementary Figure 2

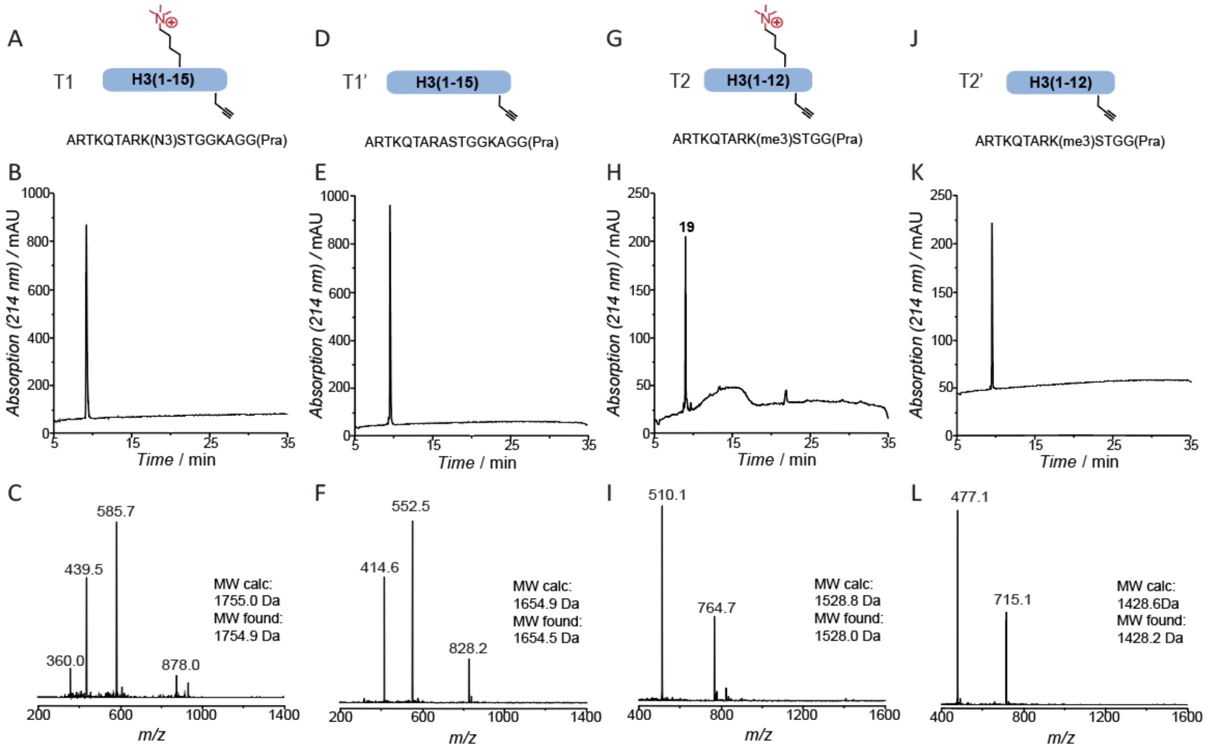


**Figure S2 – Analytics of targeting peptides. A**) Sequence of histone peptide **T1**. **B**) Analytical RP-HPLC of histone peptide **T1**. **F**) ESI-MS analysis of histone peptide **T1** (MW calculated: 1755.0 Da, MW found: 1754.9 Da). **D**) Sequence of histone control peptide **T1'**. **E**) Analytical RP-HPLC of histone control peptide **T1'**. **F**) ESI-MS analysis of histone control peptide **T1'** (MW calculated: 1654.9 Da, MW found: 1654.5 Da). **G**) Sequence of histone peptide **T2**. **H**) Analytical RP-HPLC of histone peptide **T2**. **I**) ESI-MS analysis of histone peptide **T2** (MW calculated: 1528.8 Da, MW found: 1528.0 Da). **J**) Sequence of histone control peptide **T2'**. **K**) Analytical RP-HPLC of histone control peptide **T2'**. **L**) ESI-MS analysis of histone control peptide **T2'** (MW calculated: 1428.6 Da, MW found: 1428.2 Da).

### Supplementary Figure 3

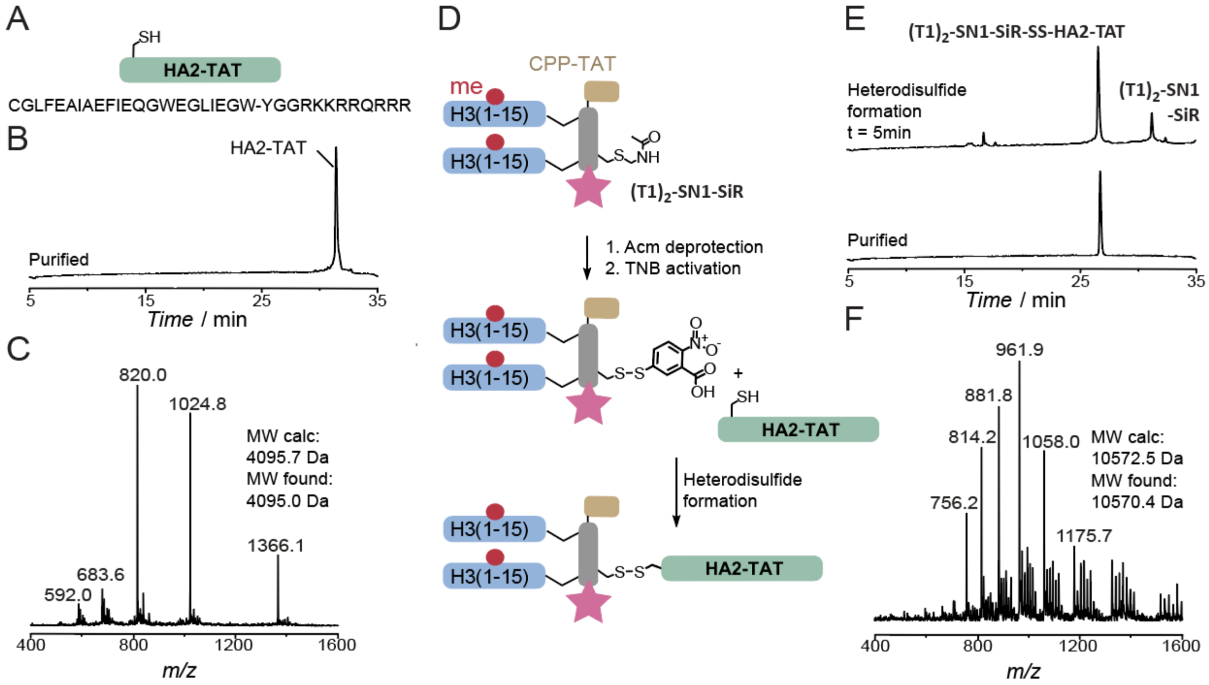


**Figure S3 – Analytics of the HA2-TAT peptide, and heterodisulfide formation. A**) Sequence of the HA2-TAT cell penetrating/endosomolytic peptide. **B**) Analytical RP-HPLC of HA2-TAT. **C**) ESI-MS analysis of HA2-TAT (MW calculated: 4095.7 Da, MW found: 4095.0 Da). **D**) Coupling of HA2-TAT to **(T1)$_2$-SN1-SiR** via a heterodisulfide linkage. **E**) Analytical RP-HPLC analysis of heterodisulfide formation between HA2-TAT and **(T1)$_2$-SN1-SiR**. **F**) ESI-MS analysis of **(T1)$_2$-SN1-SiR-SS-HA2-TAT** (MW calculated: 10572.5 Da, MW found: 10570.4 Da).

### Supplementary Figure 4

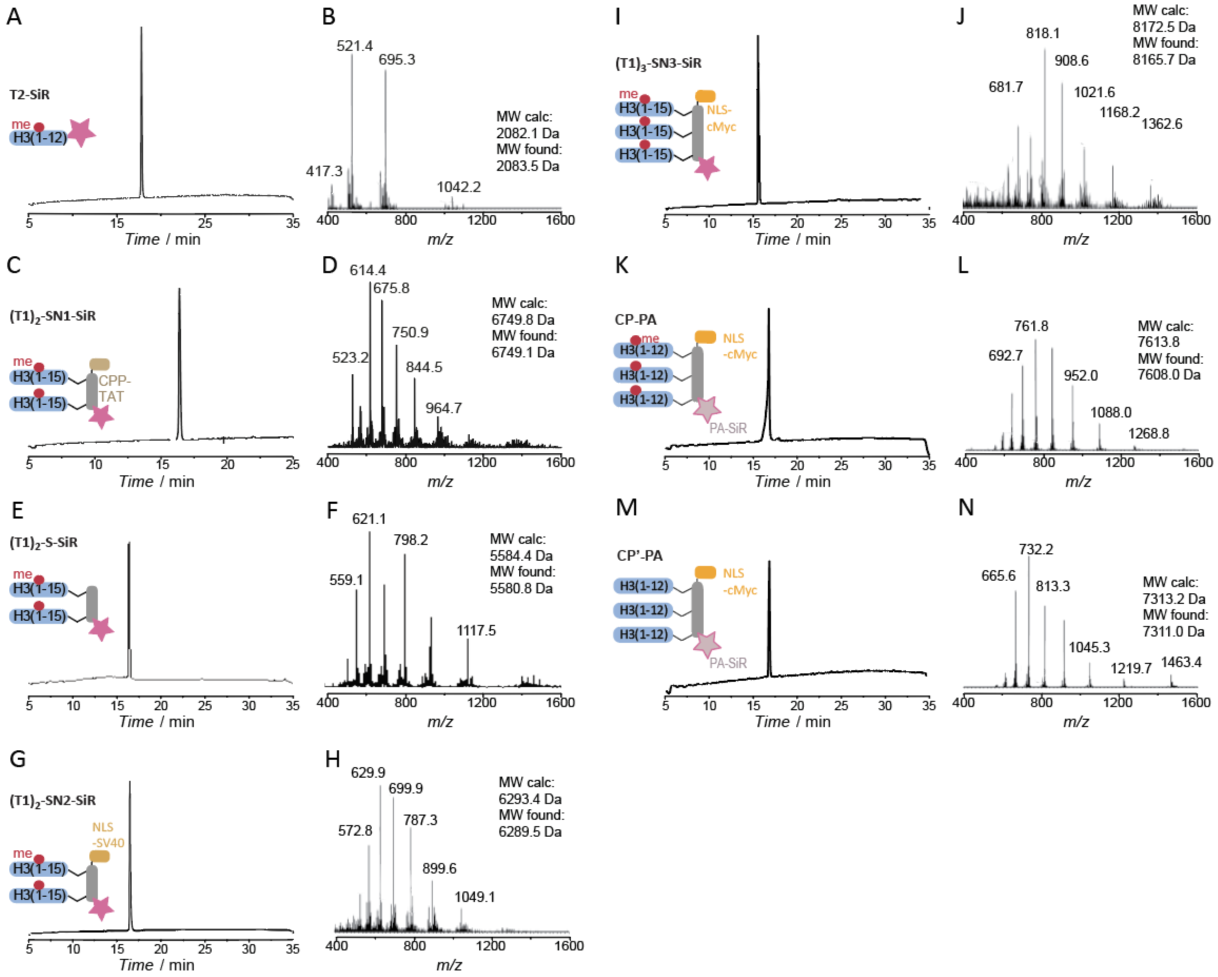


**Figure S4 - Analytics of all final peptides. A**) Analytical RP-HPLC analysis of **T2-SiR**. **B**) ESI-MS analysis of **T2-SiR** (MW calculated: 2082.1 Da, MW found: 2083.5 Da). **C**) Analytical RP-HPLC analysis of **(T1)$_2$-SN1-SiR**. **D**) ESI-MS analysis of **(T1)$_2$-SN1-SiR** (MW calculated: 6749.8 Da, MW found: 6749.1 Da). **E**) Analytical RP-HPLC analysis of **(T1)$_2$-S-SiR**. **F**) ESI-MS analysis of **(T1)$_2$-S-SiR** (MW calculated: 5584.4 Da, MW found: 5580.8 Da). **G**) Analytical RP-HPLC analysis of **(T1)$_2$-SN2-SiR**. **H**) ESI-MS analysis of **(T1)$_2$-SN2-SiR** (MW calculated: 6293.4 Da, MW found: 6289.5 Da). **I**) Analytical RP-HPLC analysis of **(T1)$_2$-SN2-SiR**. **J**) ESI-MS analysis of **(T1)$_2$-SN2-SiR** (MW calculated: 8172.5 Da, MW found: 8165.7 Da). **K**) Analytical RP-HPLC analysis of **CP-PA**. **L**) ESI-MS analysis of **CP-PA** (MW calculated: 7613.8 Da, MW found: 7608.0 Da). **M**) Analytical RP-HPLC analysis of **CP'-PA**. **N**) ESI-MS analysis of **CP'-PA** (MW calculated: 7313.2 Da, MW found: 7311.0 Da).

### Supplementary Figure 5

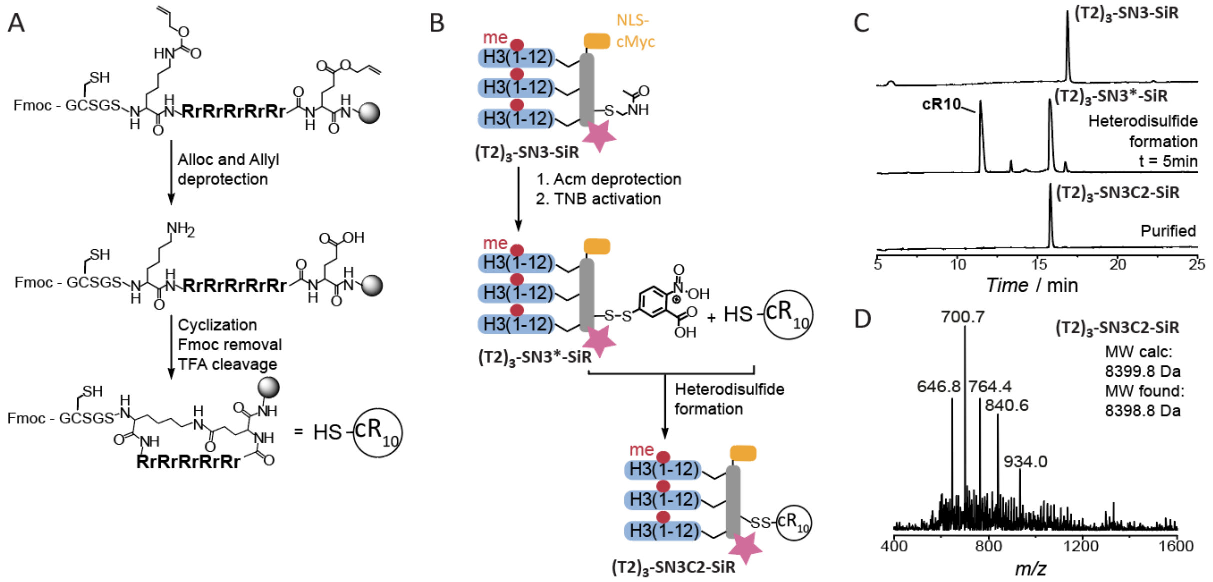


**Figure S5 – Synthesis of cR10 and coupling to peptide probes. A**) Scheme of $cR_{10}$ synthesis. **B**) Scheme of **(T2)$_3$-SN3C2-SiR** synthesis from **(T2)$_3$-SN3-SiR** and $cR_{10}$. **C**) Analytical RP-HPLC analysis of reaction progress and product **(T2)$_3$-SN3C2-SiR**. **D**) ESI-MS analysis of **(T2)$_3$-SN3C2-SiR** (MW calculated: 8399.8 Da, MW found: 8398.8 Da).

### Supplementary Figure 6

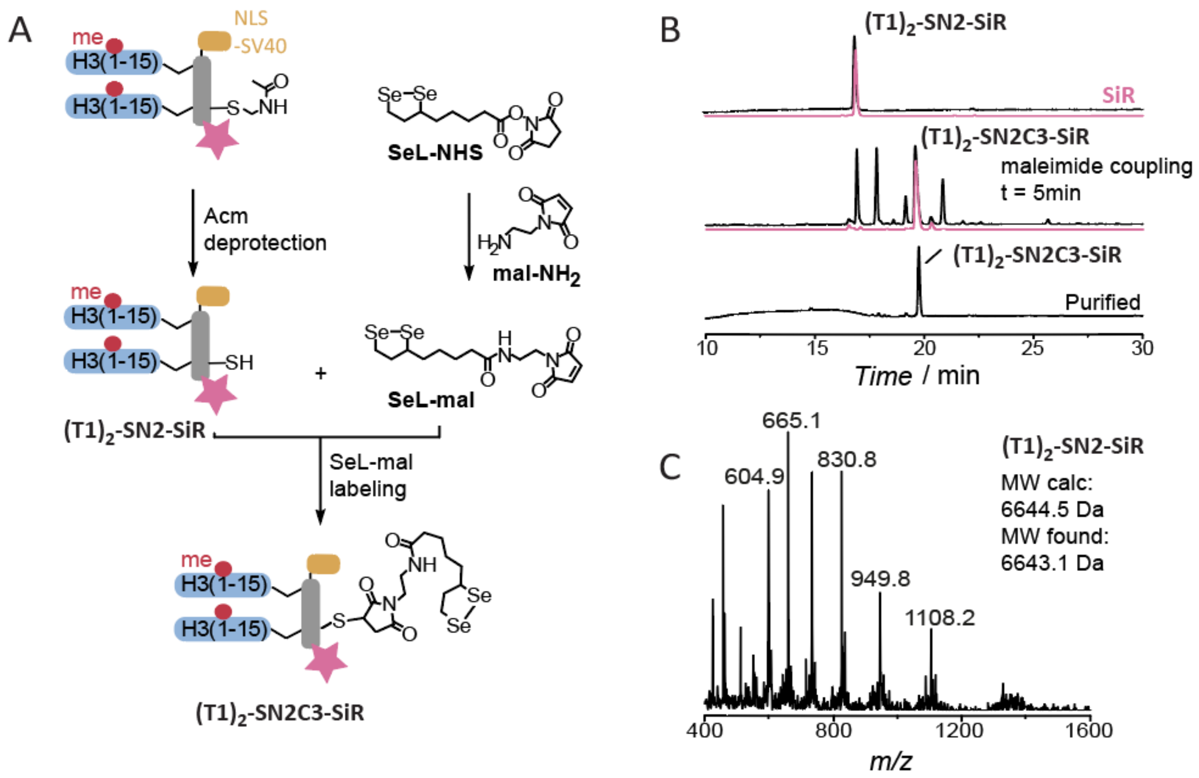


**Figure S6 – Synthesis of diselenolane probe. A**) Scheme of **(T1)$_2$-SN3C3-SiR** synthesis from **(T1)$_2$-SN3-SiR** and SeL-mal. **B**) Analytical RP-HPLC analysis of reaction progress and product **(T1)$_2$-SN3C3-SiR**. **C**) ESI-MS analysis of **(T1)$_2$-SN3C3-SiR** (MW calculated: 6644.5 Da, MW found: 6643.1 Da).

### Supplementary Figure 7

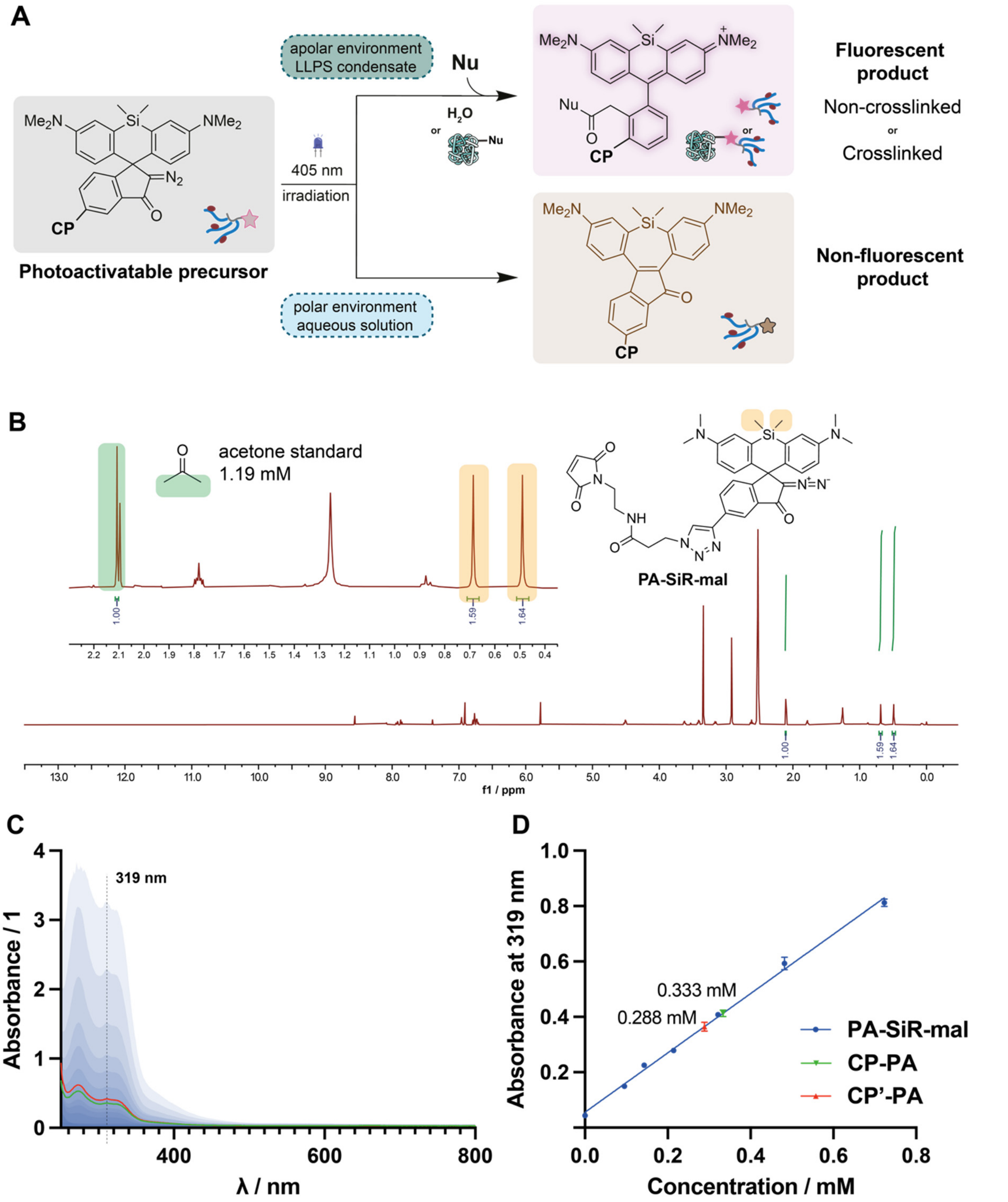


**Figure S7 – Characterization of PA-SiR**. **A**) Photochemistry of photoactivatable silicon rhodamine (PA-SiR) leading to different photoproducts. Bright photoproducts can have different mobility due to possible crosslinking. **B**) Concentration measurement of **PA-SiR-mal** using quantitative NMR with an internal standard. **C**) Spectra of **CP-PA**, **CP'-PA** conjugates and of dilution series of **PA-SiR-mal** that was used for as calibration series. No photoactivated product was detected. **D**) Concentration measurement of **CP-PA**, **CP'-PA** conjugates by absorbance at 319 nm, each datapoint represents a duplicate measurement.

**Supplementary Figure 8**

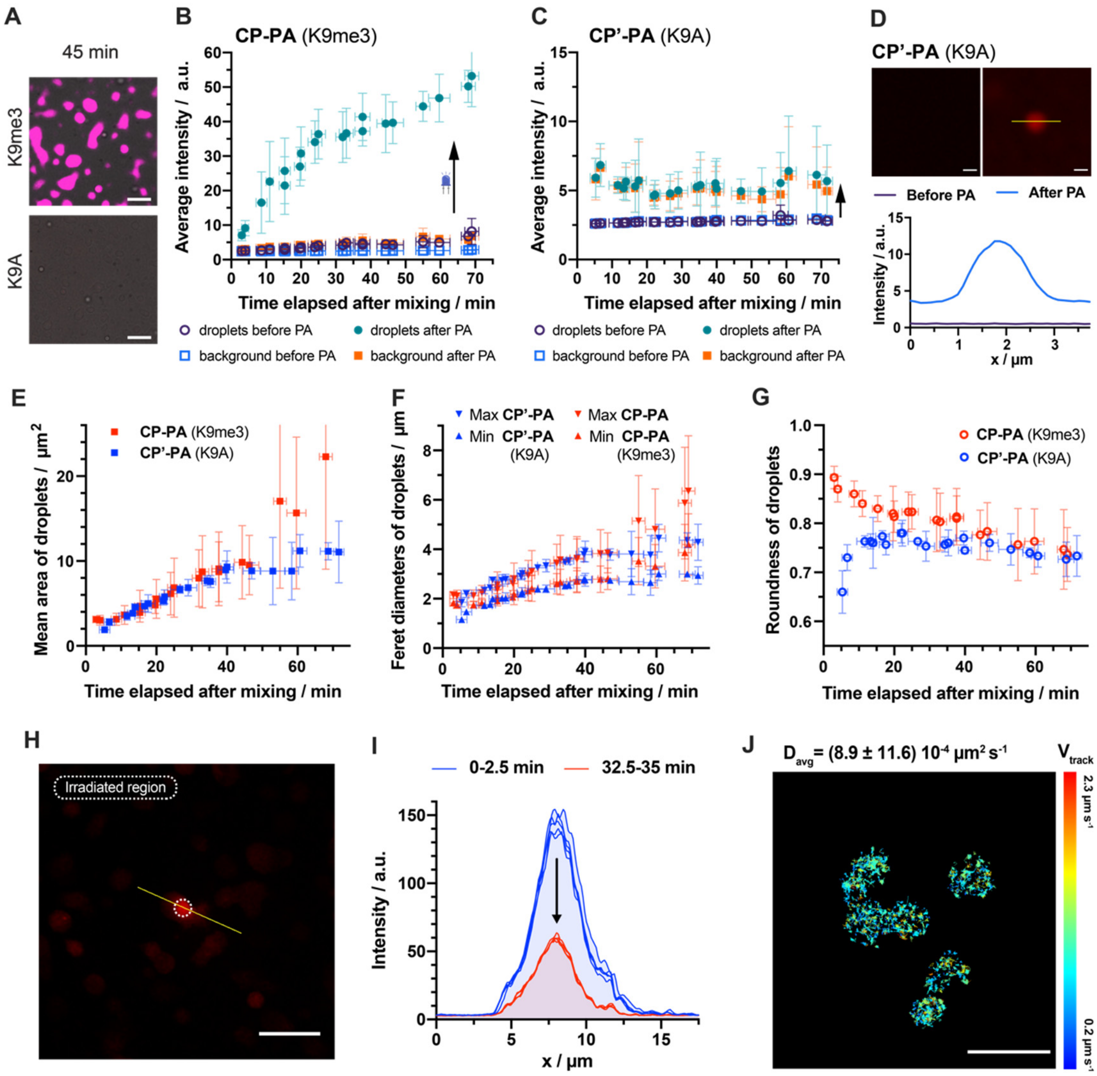


**Figure S8 – in vitro analysis of PA-droplets**. **A**) Phase-separated pHP1α droplets in presence of **CP-PA** imaged by a confocal microscope after photoactivation at 45 min (for earlier time points see **Figure 4C**). **B**) Photoactivation of **CP-PA** probe in time series with intensity measurement. Intensity was measured before and after irradiation, in droplets and at background. **C**) Photoactivation of **CP'-PA** control in time series. **D**) Photoactivation of **CP'-PA** control in droplets. Intensity cross section displays higher brightness after photoactivation than at background, proving that **CP'-PA** without enrichment in the condensate also forms bright photoproduct preferentially when irradiated. Focused 1.5 µm above glass to minimize background. **E**) Average area of droplets after mixing HP1α (30 µM) and PEG (5 $\%_w$), with **CP-PA** (100 nM). Comparison of multivalent probe **CP-PA** to control **CP'-PA**. **F**) Maximal and minimal Feret diameters of droplets to describe droplet size, after mixing. Comparing the effect of multivalent probe **CP-PA** to control **CP'-PA**. **G**) Circularity of droplets to describe the effect of multivalent probe **CP-PA** relative to control **CP'-PA**. **H**) Local irradiation of sample by 405 nm laser (10 mW, 10 µM dwell time) was done within the white circle then imaged by 638nm excitation. **I**) Irradiated sample of **H** was imaged by time lapse. Intensity profile at the beginning and end indicates no diffusion of bright photoproduct in the sample. **J**) Diffusion of single molecules inside model droplets. Tracks in are colored by mean velocity in track. Single molecules acquisitions (N=4) acquired 45 min - 2 h after mixing, average diffusion coefficient (±95% CI) value was calculated using Spot-ON assuming a single diffusing species. All error bars represent standard deviation. Scale bars: A, H = 10 µm, D, J = 1 µm.

### Supplementary Figure 9

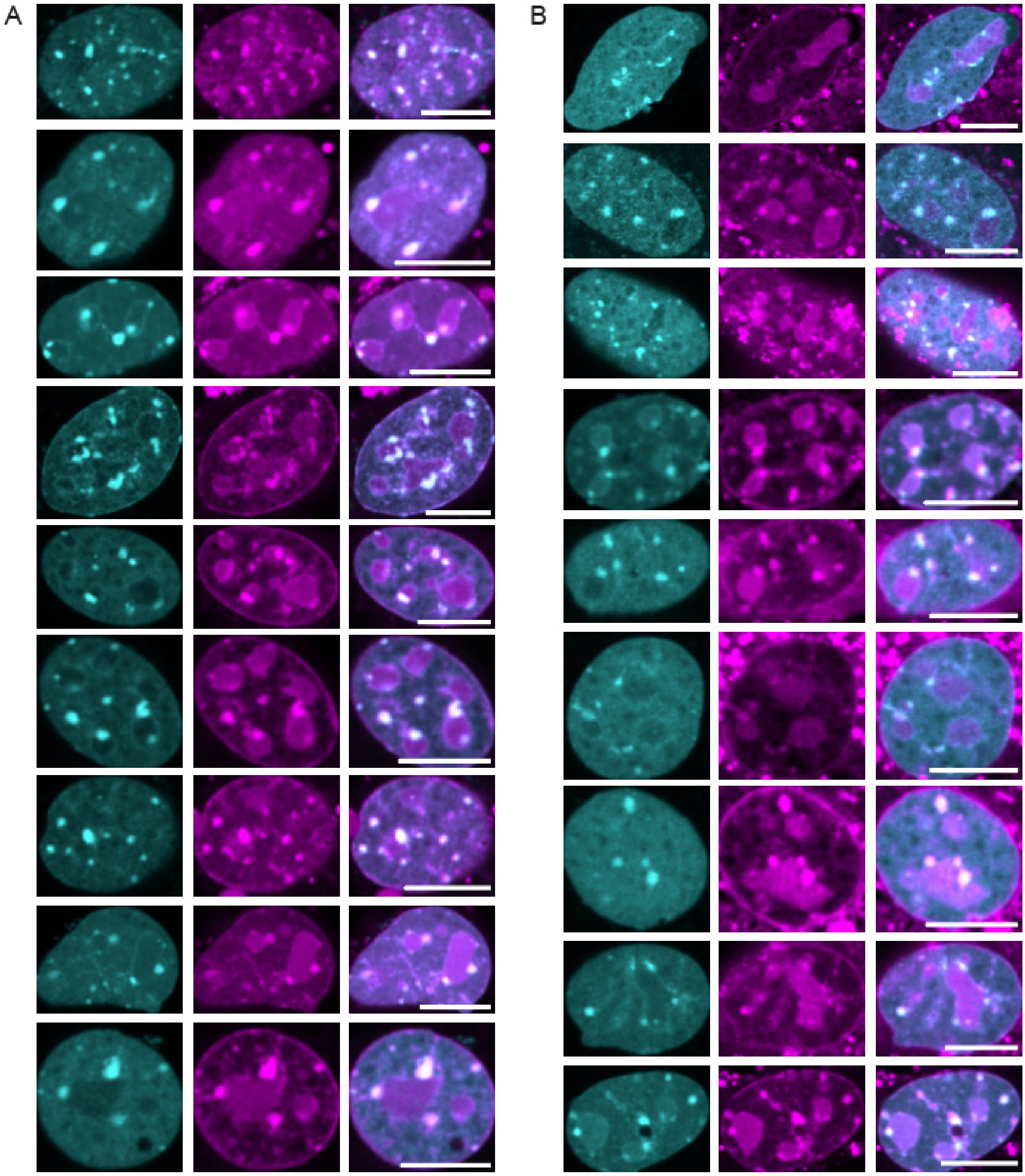


**Figure S9 – Additional images of cells of CP-SiR vs CP-PA, underlying Figure 5. A**) Heterochromatin staining using **CP-SiR**. **B**) Heterochromatin staining using **CP-PA** after photoactivation. PA-SiR after 405 nm photoactivation (magenta), HP1α-mEOs3.2 fluorescence (cyan).

### Supplementary Figure 10

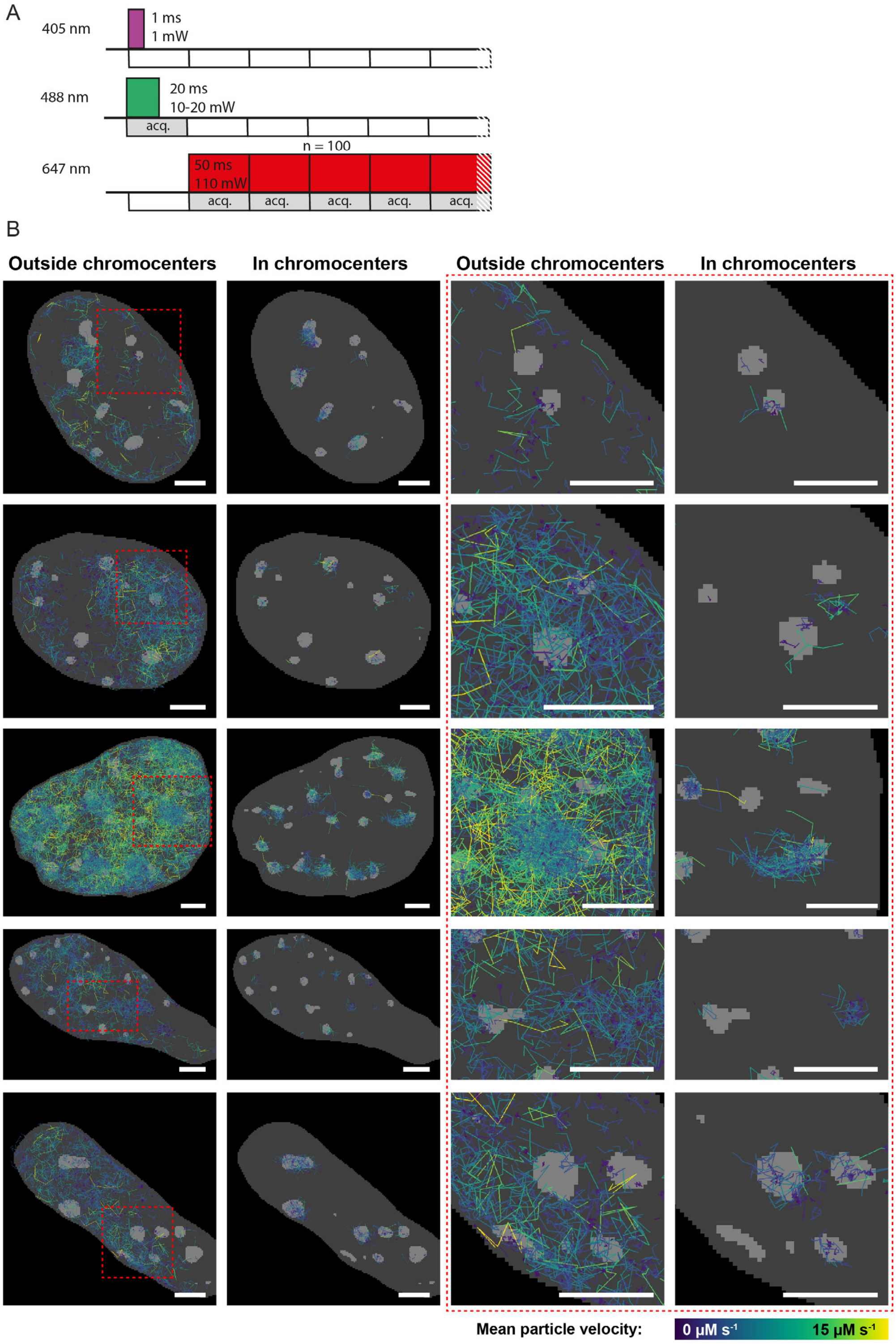


**Figure S10 – Single-molecule tracking A**) Illumination sequence timeline. **B**) Tracks of probe **CP-PA** shown overlayed with chromocenters (light grey) in the nucleus (dark grey). Tracks inside with most localization inside and outside chromocenters were grouped and are displayed in case of multiple nuclei. Tracks colored by mean velocity. Scale bar: 3 µm.

### Supplementary Figure 11

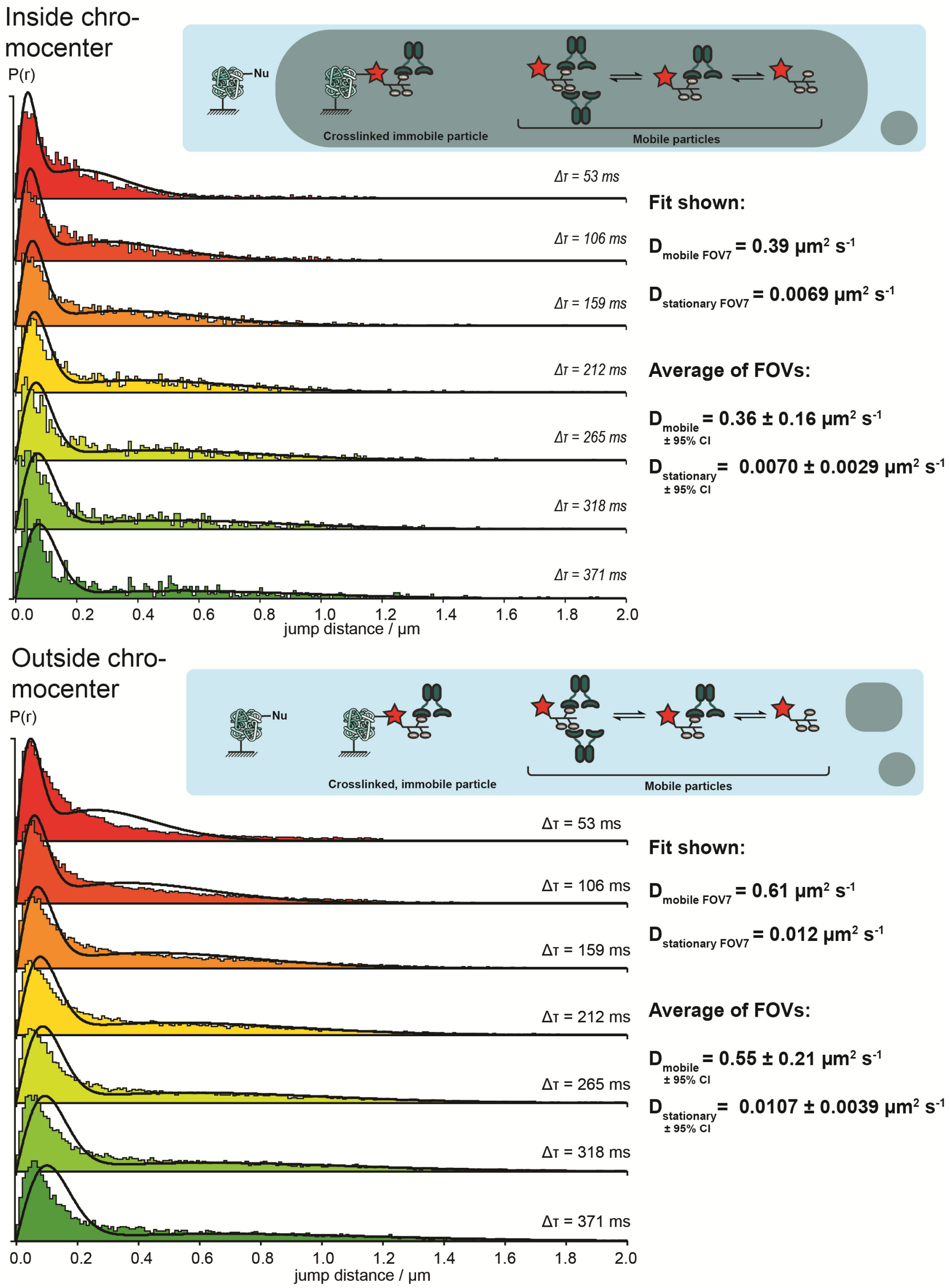


**Figure S11 – Single-molecule diffusion analysis.** Displacement histograms generated from tracks inside and outside chromocenters, and fitted assuming two diffusing species. Immobile molecules were observed both inside and outside HP1α foci, and are thought to arise form crosslinking of **CP-PA** probe upon photoactivation. There are multiple mobile species arising from the multivalent nature of **CP-PA** probe. Diffusion of these species are averaged by assuming only two diffusing species. Confidence intervals are calculated form by taking one between field of views as one measurement (N=8).

### NMR spectra

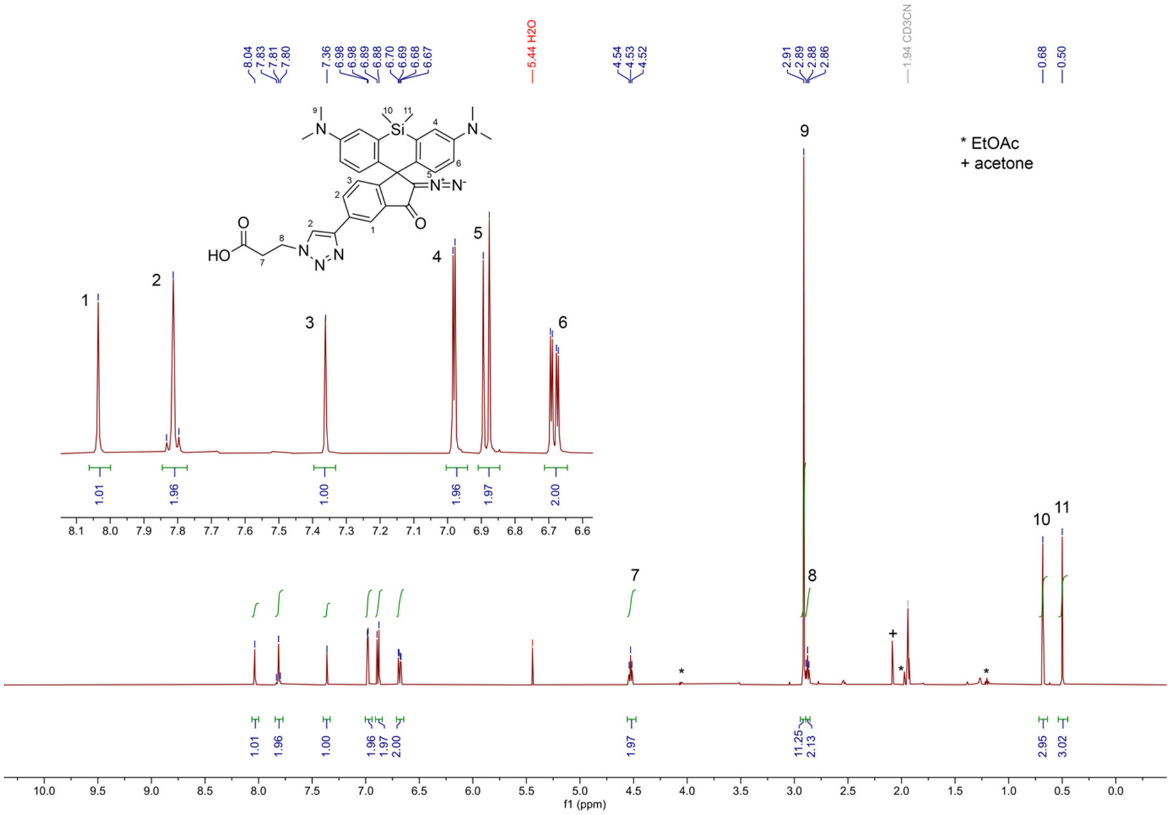

$^{1}H$ NMR (500 MHz, $CD_3CN$) spectrum of compound **PA-SiR-COOH**.

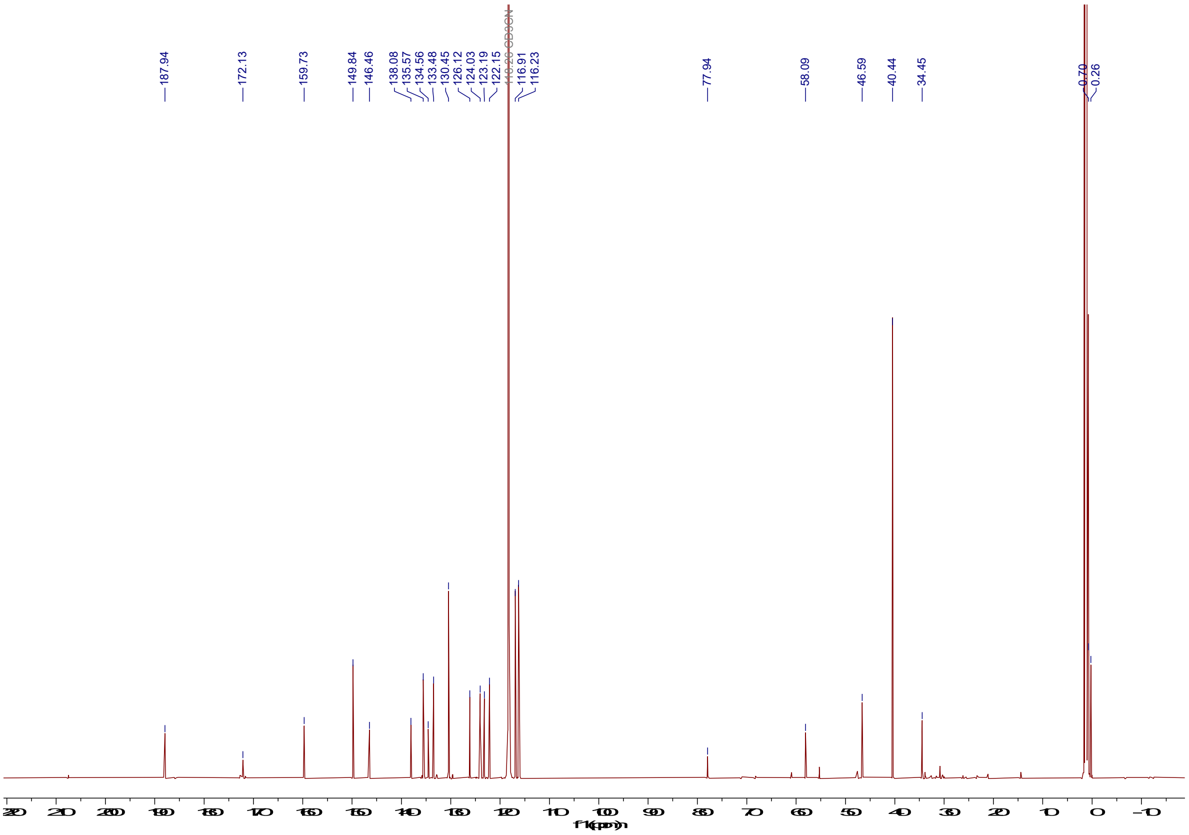

$^{13}C$ NMR (126 MHz, $CD_3CN$) spectrum of compound **PA-SiR-COOH**.

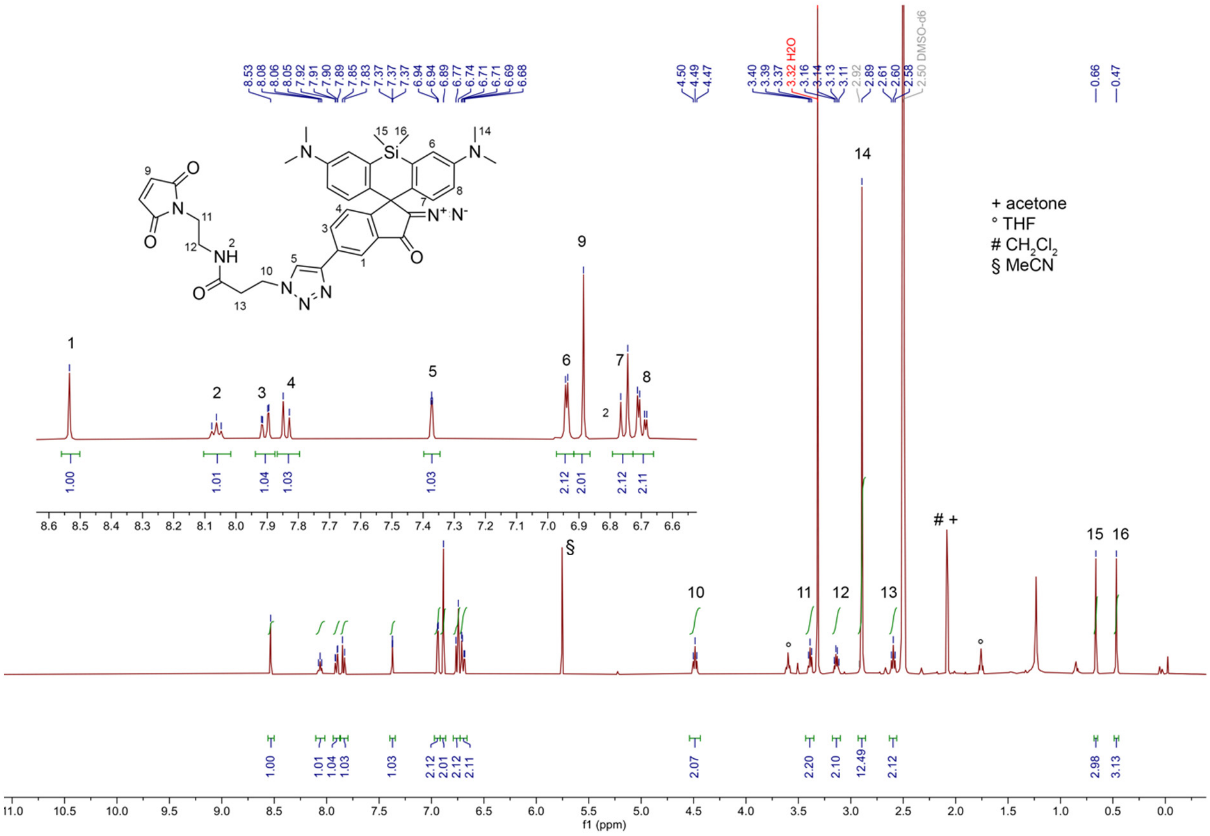


[1]H NMR (500 MHz, DMSO) spectrum of compound **PA-SiR-mal**.

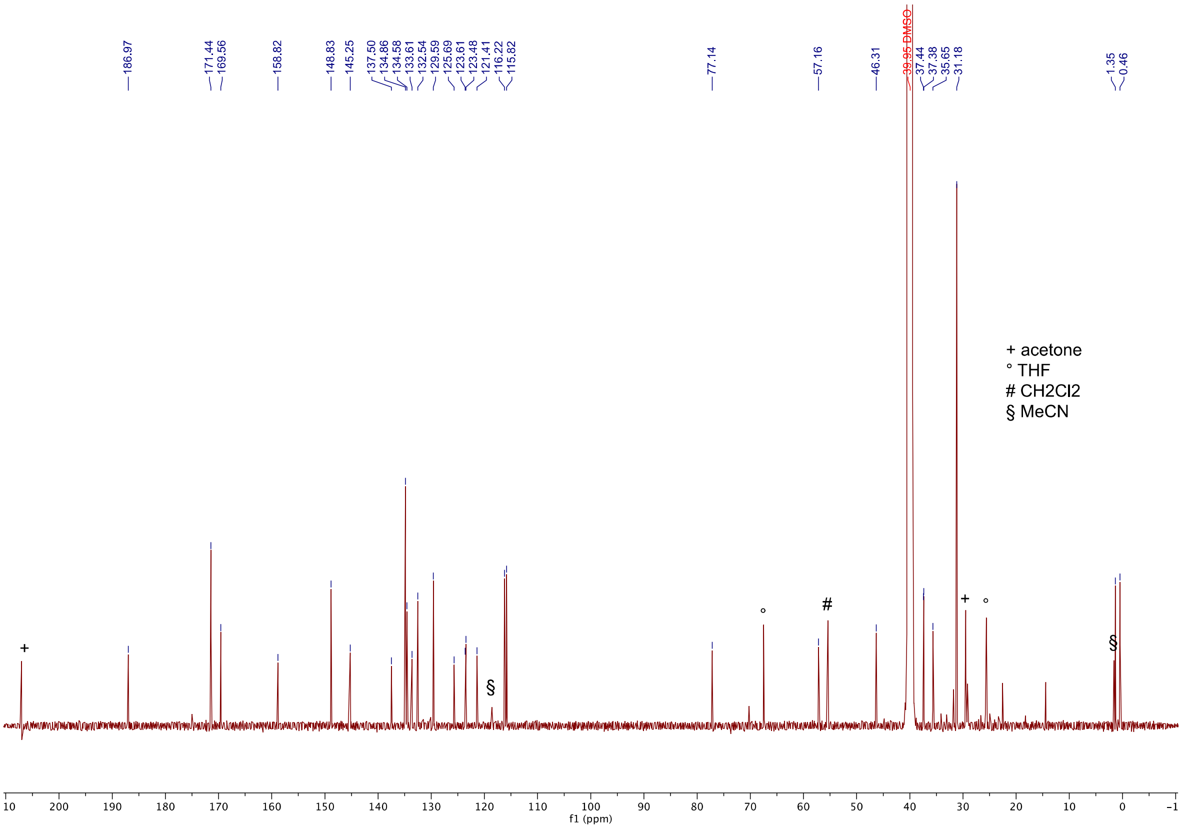


[13]C NMR (151 MHz, DMSO) spectrum of compound **PA-SiR-mal**.
